# Exploration, representation and rationalization of the conformational phase-space of N-glycans

**DOI:** 10.1101/2022.06.17.496605

**Authors:** Isabell Louise Grothaus, Giovanni Bussi, Lucio Colombi Ciacchi

## Abstract

Despite their fundamental biological relevance, structure-property relationships in *N-*glycans are fundamentally lacking, and their highly multidimensional compositional and conformational phase-spaces remain largely unexplored. The torsional flexibility of the glycosidic linkages and the ring dynamics result in wide, rugged free-energy landscapes that are difficult to sample in molecular dynamics simulations. We show that a novel enhanced-sampling scheme combining replica-exchange with solute and collective-variable tempering, enabling transitions over all relevant energy barriers, delivers converged distributions of solvated *N-*glycan conformers. Several dimensionality-reduction algorithms are compared and employed to generate conformational free-energy maps in two-dimensions. Together with an originally developed conformation-based nomenclature scheme that uniquely identify glycan conformers, our modelling procedure is applied to reveal the effect of chemical substitutions on the conformational ensemble of selected high-mannose-type and complex glycans. Moreover, the structure-prediction capabilities of two commonly used glycan force fields are assessed via the theoretical prediction of experimentally available NMR J-coupling constants. The results confirm the key role of especially *ω* and ψ torsion angles in discriminating between different conformational states, and suggest an intriguing correlation between the torsional and ring-puckering degrees of freedom that may be biologically relevant.

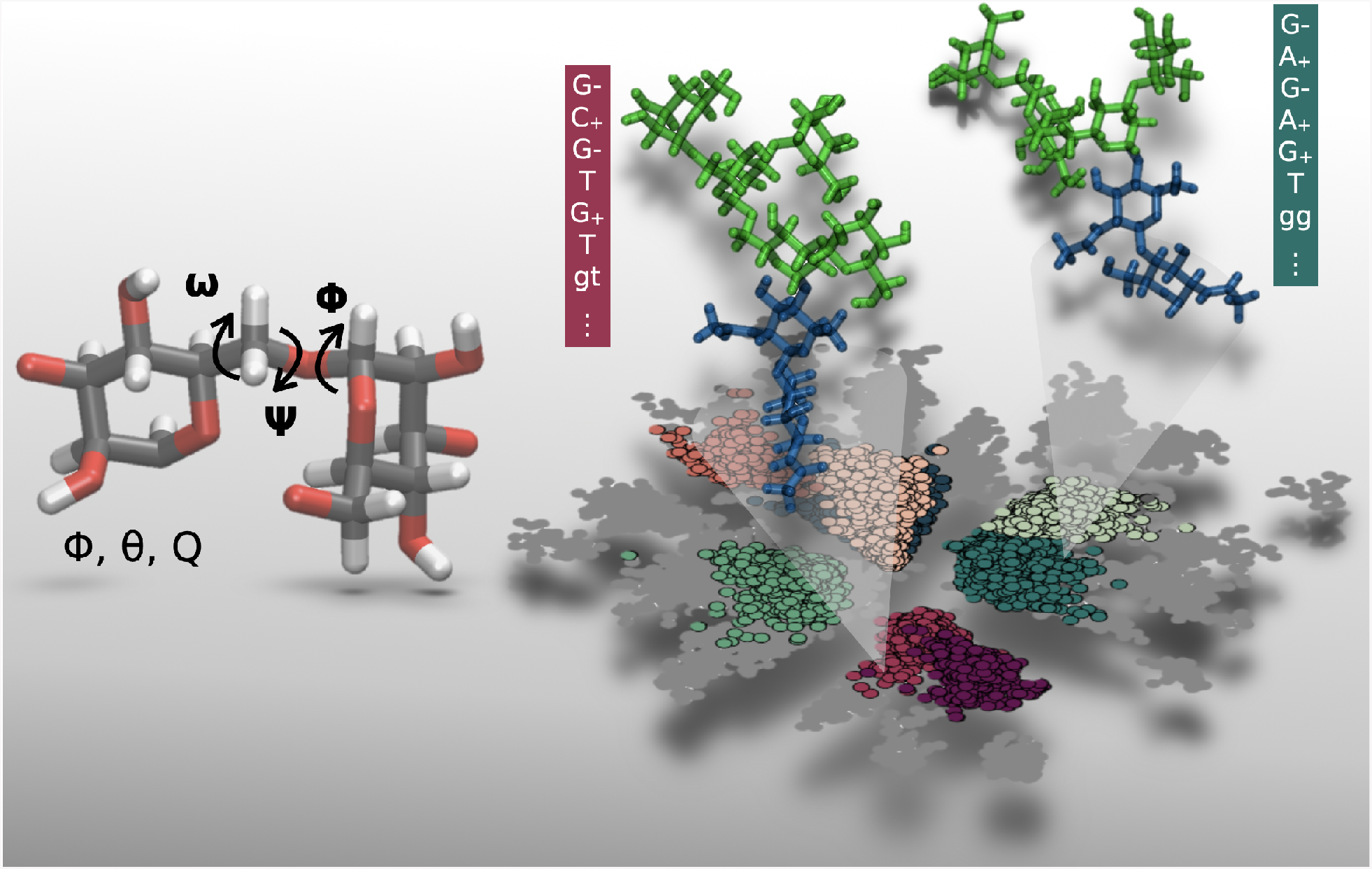

## Introduction

Biologically relevant polysaccharides, or glycans, present the largest compositional and architectural variability among all other biomolecular classes and thus harbor a rich and complex coding capability, complementing those of polypeptides and polynucleotides^1^. In fact, they do not only provide mechanical support in biological systems (e.g. as cellulose or chitin), but also act as fine-tuned encoders of information in biomolecular recognition processes. The enzyme-regulated attachment of glycans to polypeptide chains via covalent tethering to the terminal NH_2_ group of asparagine (*N-*glycosylation) is the most common post-translational modification. *N-*glycolsylation broadens the range of functionality of the underlying protein by regulating its folding, stability and function, by providing target structures for carbohydrate-binding proteins (lectins) and specific antibodies, and by mediating cell-matrix interactions as well as cell-cell recognition^2,3^. This emphasizes the need of identifying and rationalizing the rules underlying what has been named as the sugar code, namely the transmission of information mediated by glycans resulting in a specific biochemical function or signal.

Unraveling the sugar code is strongly impaired both by the lack of standard structural descriptors (such as *α*-helices and *β*-sheets in polypeptides), and by the high dynamical flexibility of glycan chains, reminiscent of the conformational variability of disordered peptides. Moreover, the non-linear, branched architecture of glycan chains and the variability of the type of linkages between the sugar monomers prevents the description of their structure in terms of few conformational variables, as done in proteins via the two-dimensional representation of all their torsional angle values in a Ramachandran plot^4^. As a first step towards a fundamental study of structure-property relationships in *N-*glycan systems, in this work we employ enhanced-sampling molecular dynamics (MD) simulations and advanced dimensionality reduction techniques to explore the high-dimensional free-energy landscape of a set of biologically relevant *N-*glycan models.

Built up of monosaccharides linked via O-glycosidic linkages, glycans harbor two main structural degrees of freedom: torsion angles and ring distortions. The *ϕ*, ψ and, in the case of 1-6 linkages, the *ω* torsion angles define the relative positions of the individual saccharide monomers (Figure 1) within each possible conformer, which are stabilized by hydrogen bonds between the hydroxyl groups of the monomers^8,9^. The distortions, or puckering, of the 6-membered saccharide rings are classified as chair (C), half-chair (H), enveloped (E), skew (S) or boat (B)^10^. These distortions can be unambiguously mapped using the spherical pucker coordinates *ϕ, θ* and *Q*, introduced by Cremer and Pople^5^. Information about the average structures adopted by *N-*glycans can be obtained by nuclear magnetic resonance (NMR) or, in some cases, by X-ray diffraction^11^. However, only atomic-scale simulations are in principle able to deliver full details of the probability distribution of all possible conformers in an *N-*glycan population, and of their dynamical behaviour^12–14^. In the case of classical MD, crucial to this regard are the accuracy of the employed force fields on one side^15,16^, and the ergodicity of the used method on the other side. In particular, the slow transitions between different conformational (rotameric) states prevent efficient phase-space sampling and convergence of conformer distributions in plain MD simulations^17,18^.

**Figure 1:**
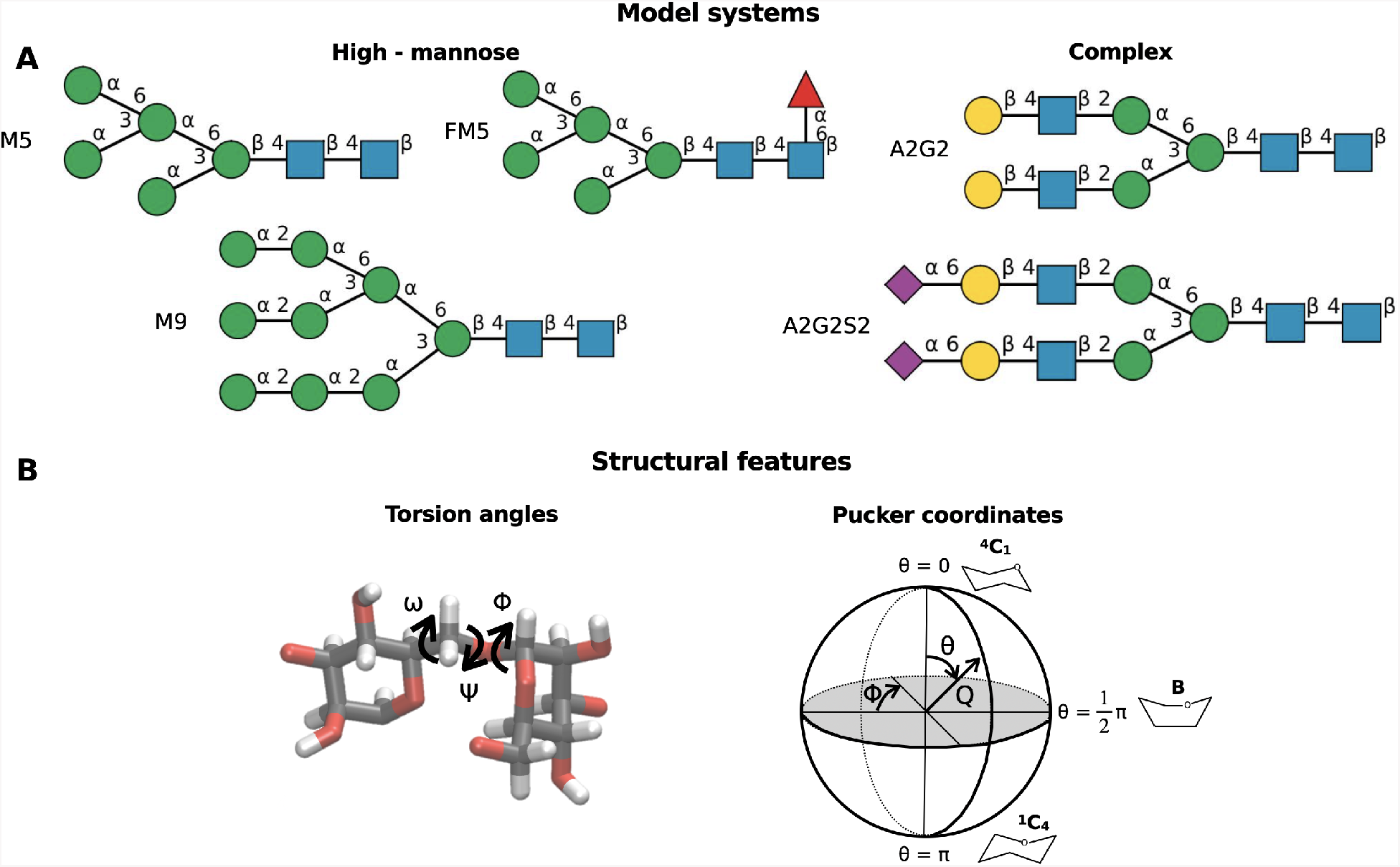
Top: Model systems employed in this study, namely three high-mannose type *N-*glycans (M5, FM5, M9) and two complex *N-*glycans (A2G2, A2G2S2). Bottom: Representative atomistic details are described for a torsion angle along a 1-6 O-glycosidic linkage as well as for the pucker coordinates according to the Cremer-Pople representation^5^. Monosaccharide symbols are in congruence with the nomenclature of the Consortium for Functional Glycomics^6^, using the Oxford notation for *N-*glycan names. The models were drawn using DrawGlycan^7^.

For this reason, various enhanced-sampling MD techniques have been used to facilitate the crossing of the relevant energy barriers and accelerate the transition probabilities. These include replica-exchange MD (REMD)^19^, Hamiltonian replica-exchange MD (H-REMD) with solute scaling (REST2)^20^, well-tempered metadynamics (WT-MetaD)^21^, Umbrella Sampling (US)^22^ and their different combinations. On the one hand, methods based on bias potentials applied to specific collective variables (CVs), such was WT-MetaD, have so far focused on only few specific torsion angles (e.g. *ω*)^18^, not giving justice to the structural complexity of *N-*glycans with multiple branches ^18,23–25^. On the other hand, CV-independent methods such as REMD do not guarantee complete phase-space sampling^18,23,26^, and require elaborate pre-calculations when used together with additional bias potentials^24,27^. To overcome these difficulties, in this study we combine REST2 with the recently introduced replica-exchange with collective-variable tempering (RECT) algorithm^28^ (Figure 2). RECT enhances the transition probability between conformers separated by high energy barriers by means of one-dimensional bias potentials applied to a large number of selected degrees of freedom, e.g., all torsion angles, in a MetaD framework. Simultaneously, REST2 samples all other degrees of freedom via a solute-scaling approach which requires only a low number of replicas.

**Figure 2:**
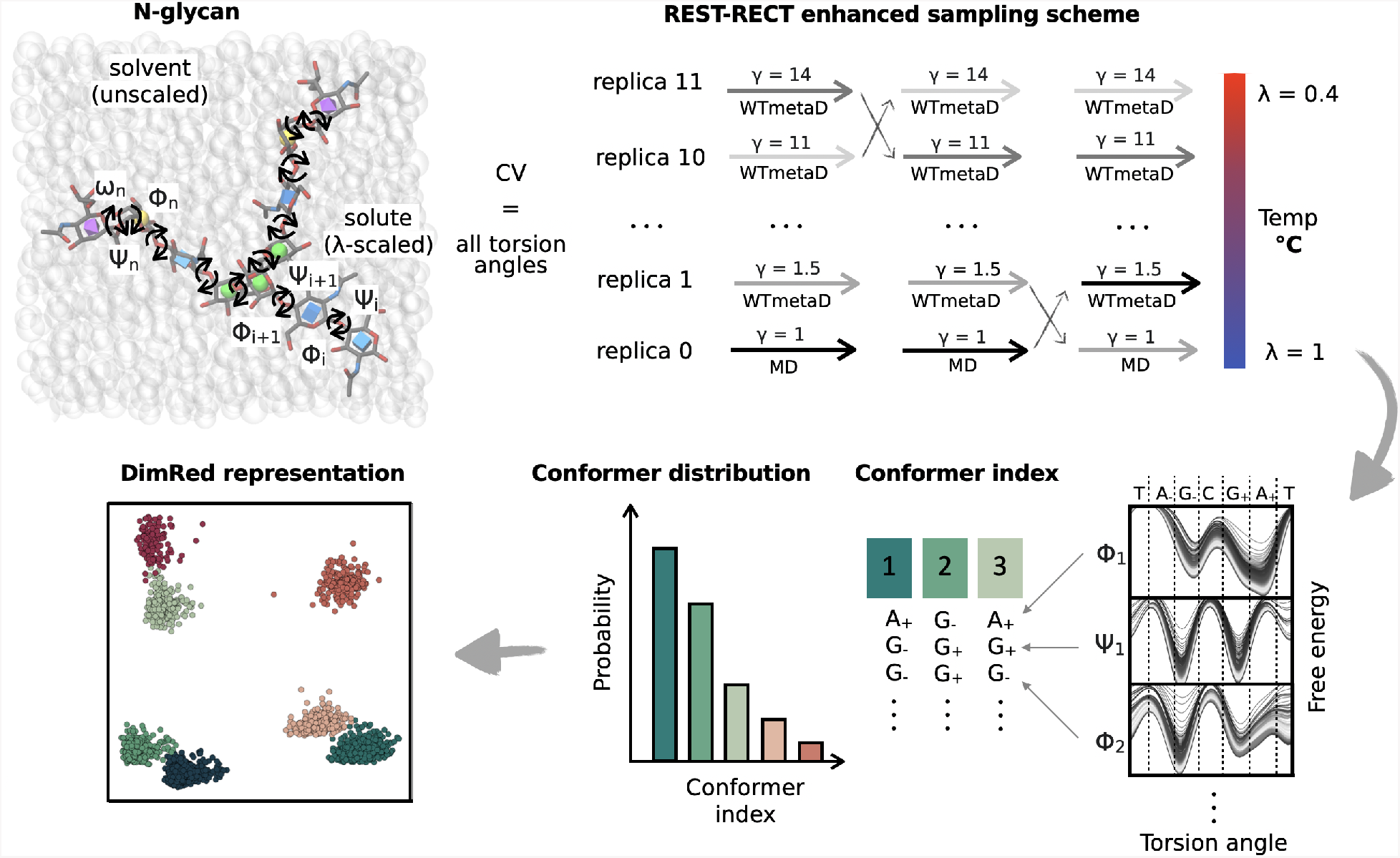
Schematic overview of the approach followed in this study. Free *N-*glycans in solution are simulated employing the enhanced-sampling method REST-RECT to accelerate transitions over barriers for all torsion angles and pucker coordinates. Conformer strings are then constructed based on the free energy landscape of each torsion angle. Individual conformers are grouped together according to these strings, and conformer distributions constructed from the simulated trajectories. Low-dimensional representations of the conformer clusters are finally generated using dimensionality-reduction methods.

Due to the large number of CVs used to define the conformational states of an *N-*glycan, structural analyses require the rationalization of a high-dimensional vector space. Dimensionality reduction techniques are thus required for an efficient graphical representation of the glycan conformational space and of its associated free-energy landscape. Such reduction should be performed in a way that best differentiates among the most probable conformers, possibly revealing mutual functional dependencies and hidden correlations among the many CVs. Diverse strategies have been developed in recent years to map a high-dimensional feature matrix **X** onto a low-dimensional latent-space matrix **T**, and their applicability mainly depends on the underlying data structure^29^. For instance, Principle Component Analysis (PCA)^30^ projects the data onto the linear eigenvector space defined by the *k* largest eigenvalues obtained by diagonalization of the covariance matrix of **X**. Therefore, the top *k* eigenvectors (the principal components) represent the maximum data variance and are used as the axes of two-dimensional or three-dimensional graphs representing the entire data space. In contrast, Diffusion Map^31^ is a non-linear method, where the connectivity (or diffusion distance) between individual data points (in our case, individual *N-*glycan conformers) is quantified by the likelihood of transitioning from one to the other, as expressed with the help of a diffusion kernel function. Data points of **X** are projected onto a two-dimensional matrix **T** in such a way that the diffusion distances in the high dimensional feature space can be approximated by Euclidean distances between points in the reduced space, ensuring preservation of the local vector-space structure (so-called isometric embedding). The non-linear Sketch-map algorithm^32,33^ also ensures isometric embedding, but is based on a different approach named multidimensional scaling (MDS)^34^. The mutual distance of data points in **X** is conserved in **T** by the application of a sigmoid function focusing on the reproduction of intermediate distances, rather than far-away distances which are dominated by the topology of the high-dimensional space. Finally, beside these unsupervised techniques, there are also supervised algorithms, which make use of information included in an additional property matrix **Y**. A recent example is the kernel principal covariates regression model (kPcovR)^35^, which combines the robust methods of linear regression and PCA.

In this study we investigate the ability of all these four dimensionality reduction techniques to effectively measure the similarity among *N-*glycan conformers, and partition them into meaningful subgroups (clusters) for further analysis. The proposed simulation and analysis scheme (Figure 2) is applied to different *N-*glycans (Figure 1) that are dominant in human plasma samples and therefore represent valuable model systems^36^. Our aim is to reveal how differences in the chemical composition of *N-*glycans impact the probability distribution of their conformers. In this way, the *N-*glycan heterogeneity can be better correlated with their biological functions. Additionally, we assess the performance of two force fields, namely CHARMM36m^37^ and GLYCAM06j^38^, in predicting correct torsion angle and pucker distributions by comparison of our computational results with experimental NMR data.

## Methods

### Simulation setup

Five different *N-*glycans were investigated in this study, namely M5, FM5, M9, A2G2S2 and A2G2 (Figure 1). Their three-dimensional structures were constructed using the CHARMM-GUI Glycan Modeller, based on averaged structures from the Glycan Fragment Database^39–42^. The *N-*glycans were solvated with a 15 Å thick water layer in a cubic box. Two sodium counter-ions were added to compensate for the two net negative charges of A2G2S2.

### MD simulations

MD simulations were performed with the GROMACS code, version 2018.4^43^, patched with the PLUMED package, version 2.6^44^. Either the CHARMM36m^37,45,46^ or the GLYCAM06j^38^ force field were used for the *N-*glycans molecules in combination with the CHARMM-modified TIP3P water model (mTIP3P)^47^ or the standard version (sTIP3P)^48^, respectively. In the case of the CHARMM36m force field, the recent correction to a previous faulty implementation affecting in particular the ring inversion properties of Neu5Ac has been applied in all simulations^49^. The leap-frog algorithm was used as an integrator with a 2 fs time step and the LINCS algorithm^50^ was employed to constrain bonds connected to hydrogen atoms. Temperature control was realized via velocity rescaling^51^ using a time constant of 0.1 ps, setting a reference temperature of 310.15 K. The pressure was set to 1 bar with a compressibility of 4.5 × 10^−5^ bar^−1^, and kept constant via the Parrinello-Rahman barostat with a time constant of 5 ps. The Verlet list scheme^52^ was employed with a neighbor list updated every 80 steps. The calculation of electrostatic interactions was done with the Particle Mesh Ewald (PME)^53^ method using a cut-off distance of 1.2 nm for the real space contribution. The following steps were performed to equilibrate the systems. First, an energy minimization of water and ions was performed using the steepest-descent algorithm with a tolerance of 1000 kJ mol^−1^ nm^−1^, restraining the *N-*glycan atoms. Then, the solvent was equilibrated in one NVT and one NPT run, each lasting 1 ns, with restrained *N-*glycans. After that, a second energy minimization of all atoms was performed with no constraints, with the same parameters as before. Finally, unrestrained NVT and NPT equilibration runs were performed, lasting 1 ns and 100 ns, respectively. For each simulated system, two different starting conformers, named s1 and s2, were generated by setting the *ω* angle of the **6**– branch (see below for the definition of the branch labelling) either in a gt (for s1) or a gg (for s2) conformation. Separate simulations starting with the two initial conformations s1 and s2 were performed to validate the convergence of the conformer distributions predicted by the different simulation techniques employed. For plain MD simulations, each production run lasted 6 *µ*s, recording frames every 8 ps.

### Enhanced-sampling simulations

Enhanced-sampling MD simulations were performed with a combination of the REST2 Replica Exchange method^20,54^ and with the RECT^28^ method based on WT-MetaD^21^ (henceforth referred to as REST-RECT). In each simulation, the whole *N-*glycan was defined as the REST2 solute, whose temperature was scaled in *i* replicas by means of scaling factors *λ*_*i*_ acting on the long range electrostatics, the Lennard-Jones interactions, as well as the dihedral angles. For non-neutral systems a neutralizing background was automatically added via GROMACS. We used 12 replicas and a geometric progression of *λ*_*i*_ values equal to 1, 1, 0.92, 0.84, 0.77, 0.71, 0.65, 0.60, 0.55, 0.50, 0.46, 0.42, spanning an effective temperature ladder from 310.15 K to 800.00 K. Note that both the ground replica (*i* = 0) and the first replica (*i* = 1) were at the same ground temperature *T*_0_ = 310.15 K, for convenience of the RECT implementation and analysis (see below). Water and ions were always kept at the ground temperature. Replica exchanges were attempted every 400 steps, following a Metropolis-Hastings acceptance criterion. In the RECT part, all *n* torsion angles of the simulated glycan were defined as CVs and biased simultaneously by *n* one-dimensional potentials in each replica *i. n* amounted to 14, 17, 24, 17 and 23 in the M5, FM5, M9, A2G2 and A2G2S2 *N-*glycans, respectively. Torsion angles were defined as *ϕ* = O5′–C1′–O*x*–C*x*, ψ = C1′–O*x*–C*x*–C(*x*–1) and *ω* = O6–C6–C5–O5, with *x* being the carbon number of the linkage at the non-reducing end. An exception are the 2–6 angles between Gal and Neu5Ac, which are defined as *ϕ* = O6′–C2′–O6–C6, ψ = C2′–O6–C6–C5 and *ω* = O6–C6–C5–O5. The *i* replicas were biased with bias factors *γ*_*i*_ following a geometric progression of values equal to 1, 1.2, 1.46, 1.82, 2.3, 2.94, 3.78, 4.89, 6.34, 8.23, 10.7, 14. Gaussian hills were deposited at time intervals of *τ*_*G*_ = 1 ps, with a width of 0.35 rad and a height corresponding to *h*_*i*_ = (*k*_*B*_Δ*T*_*i*_*/τ*) *× τ*_*G*_, where *k*_*B*_ is the Boltzmann constant, Δ*T*_*i*_ = *T*_0_(*γ*_*i*_ − 1) the boosting temperature, *τ* = 4 ps the characteristic time for the bias evolution in the RECT part. The geometric progressions of *λ*_*i*_ and *γ*_*i*_ ensured sufficient overlaps of the potential energy distributions at all temperatures, resulting in uniform round trip times for the different replicas (Figure S3, S5). We note that the ground replica was fully unbiased (*λ*_0_ = 1, *γ*_0_ = 1), allowing for an unbiased statistical distributions of frames, which could be trivially used in all subsequent analyses. The first replica was biased, but kept at the ground temperature (*λ*_1_ = 1, *γ*_1_ = 1.2) to ensure sufficient overlaps between the first two replicas. The inclusion of higher-order replicas in the analyses requires the application of the Weighted Histogram Analysis Method (WHAM)^55^. However, including replicas up to *i* = 4 in the analyses did not result in a more effective sampling. Higher replicas should not enter the analyses because of the expected low ensemble overlap with the reference replica.

### Conformer string generation

*N-*glycans are multi-branched structures, characterized by the specific linkages between saccharides monomers. Each glycosidic linkage gives rise to at least two torsion angles (*ϕ* and ψ), where 1–6 and 2–6 linkages harbor an additional torsion angle *ω*. Based on these structural characteristics, we constructed an unambiguous labeling scheme to distinguish different conformers of the same *N-*glycan (see Figure 3 for examples related to M5 and M9 glycans). The scheme is also applicable to other glycans, independently of their size, number or type of branches and amount of substitutions such as fucosylation. Each conformer is identified by a digit string of length *n* equal to the number of torsion angles in the glycan. The string begins at the free reducing end of the glycan, which can be linked to the asparagine residue of proteins by oligosaccharyltransferase enzymes. *N-*glycans always start with a *β* 1 − 4-linked GlcNAc dimer, followed by a mannose residue (compare Figure S1). For each linkage, the string reports digits assigned to *ϕ*, ψ and *ω* (if applicable), in this order. In correspondence of a junction (leading e.g. to an *α* 1–6 and a *α* 1–3 branch after the first mannose), a string separator is introduced, labelled according to the C atom at the branch origin (e.g. **6**– for 1–6 linkages). The string continues first along the branch of the higher C atom (6 in our case) until reaching the terminal residue, prior to returning to the last junction and following the branch of the next-lower C atom (3 in our case). Additional modifications like the attachment of fucose residues or bisecting GlcNAc residues are included after all other branches are assigned. Since our models have only one or two junctions, the separators of primary branches are labelled with bold numbers (**6**– or **3**–), for clarity. The string digits indicate in which intervals of values the torsion angle lies, following the IUPAC nomenclature for dihedrals ^56^. Namely, the digits for *ϕ* and ψ and the corresponding interval of radian values are C = [−0.52, +0.52), G_+_ = [+0.52, +1.57), A_+_ = [+1.57, +2.62), T = [+2.62, *π*] or [−*π*, −2.62), A_−_ = [−2.62, −1.57), G_−_ = [−1.57, −0.52). The digits for *ω* are gg = [−2.62, 0), gt = [0, 2.62), and tg = [2.62, *π*] or [−*π*, −2.62). The assignment of each torsion angle to a given interval is performed in the following way. First, the free-energy profile associated with rotation along the torsion angle is calculated from all frames of the MD trajectories (either plain MD or REST-RECT MD). The position of the free-energy minima are then labelled according to the nomenclature above. All angles belonging to the same free-energy basin (around a minimum between the two neighboring maxima) are finally labelled equally to the minimum of their basin.

**Figure 3:**
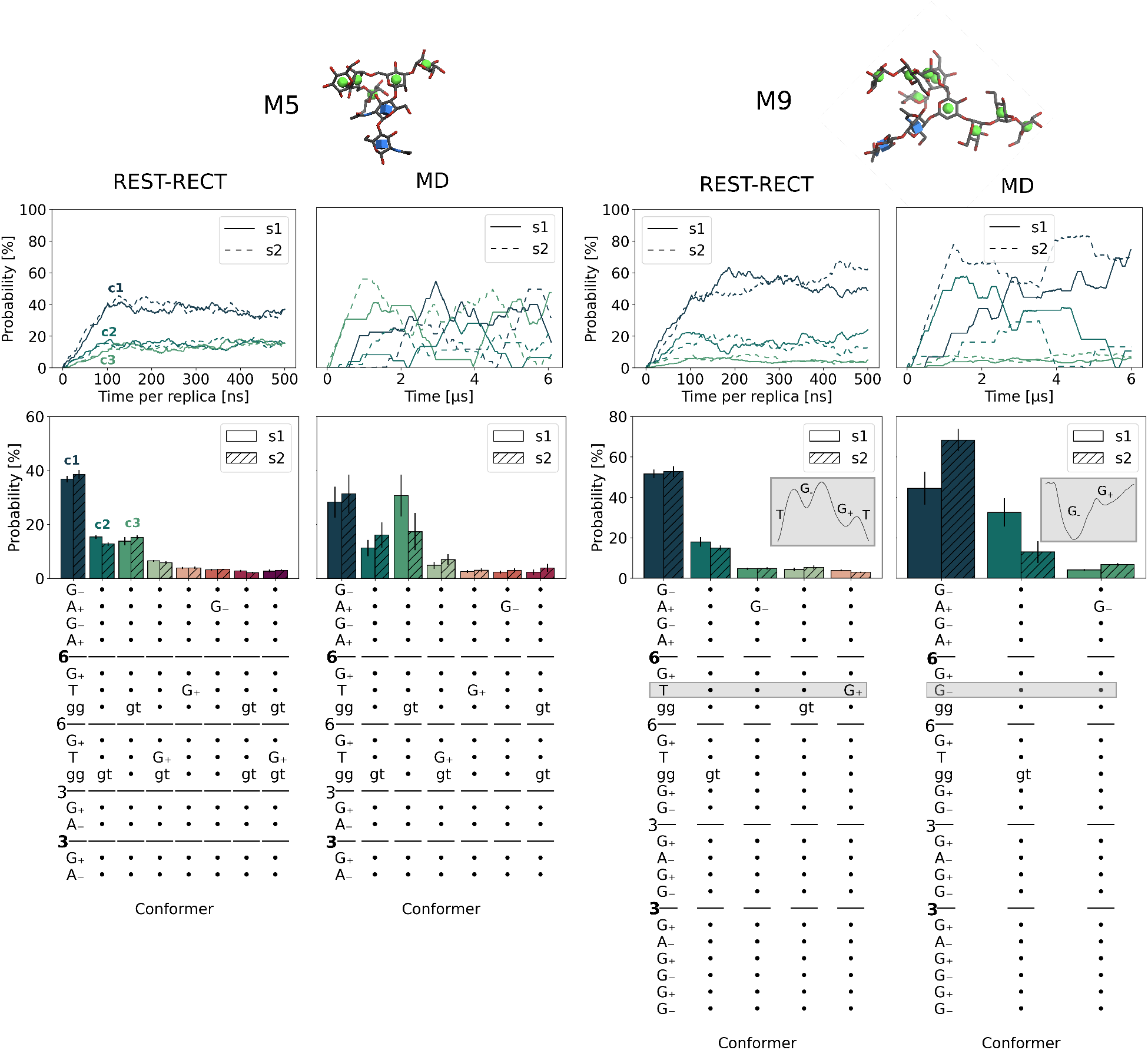
Comparison of REST-RECT with plain MD simulations for systems M5 and M9 using the CHARMM36m force field. The upper panel includes atomistic structures of each glycan, visualized with the 3D-SNFG tool of VMD^65^. The middle panel shows the moving average for the three most populated conformer clusters using a window size of 100 ns (REST-RECT) and 1.2 *µ*s (MD), corresponding to the same sampling time. Two separate simulations were performed with differing initial starting configurations (s1 and s2, see Methods). The lower panel reports the resulting conformer distributions. The conformer string is given on the x-axis, where each digit stands for a torsion angle, the letter representing the occupied free-energy minima (see Methods). Dots are used instead of letters when no change could be observed in comparison with the most populated conformer cluster. The gray boxes highlight a key conformational difference between REST-RECT and MD simulations, with a depicted free energy landscape of the corresponding angle (inserts). Only conformers with a probability higher than 2.5 % are plotted.

### Probability distributions

In all simulations, torsion angle values were recorded for each frame and converted into conformer strings. Histograms were constructed using the individual conformers as bins. In order to assess statistics, block averaging was performed, separating the data set in evenly distributed blocks. The average of all blocks 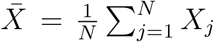 was calculated over *N* = 10 blocks, where *X*_*j*_ is the average calculated within each *j*th block. Error bars were calculated as standard deviations of the sampling distributions (standard error of the mean): 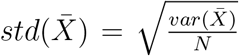, with the variance of the sampling distributions 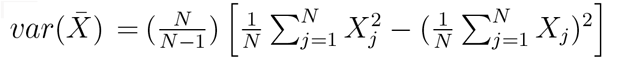.

### Dimensionality reduction

Four different dimensionality reduction techniques were compared in this study. We always included 31250 data points (*n*_*samples*_), with the different torsion angles of each *N-*glycan defined as features (*n*_*features*_), resulting in a feature matrix **X** with shape *n*_*samples*_ *× n*_*features*_. In order to account for the periodicity of the torsion angles, their sin and cos values were used in the feature matrix **X** instead of the torsion angles in all cases except for the Sketch map.

### Principal component analysis

The PCA calculations were performed with the scikit-learn package^57^. Whenever two different feature matrices **X** were compared to each other, e.g. stemming from simulations performed with two different force fields, the corresponding data sets were concatenated prior to PCA calculation. Free energy differences (Δ*G*) along the principal components 1 and 2 defining the low-dimensional latent-space matrix **T** were calculated by constructing two-dimensional histograms with 35 bins and converting the histogram probabilities *P* according to Δ*G* = −*k*_*B*_*T* ln(*P*).

### Diffusion maps

Diffusion maps were computed based on the same algorithm as described by Bottaro et al.^58^, using the Gaussian kernel 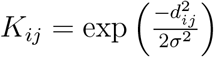, where *d*_ij_ represents the pairwise distance matrix between sin or cos of the torsion angles values of conformers *x*_*i*_ and *x*_*j*_. The parameter *σ*, defining the size of the neighborhood including similar conformational structures, was set equal to 1.7. We note that the used algorithm is different from the original implementation^31^ in that the transition matrix is simultaneously normalized and made symmetric by means of an iterative procedure^58^. This yields results that are equivalent to the recently introduced bi-stochastic kernel method^59^.

### Sketch-map

Sketch-maps were calculated with the DimRed module of PLUMED (version 2.6). The matrix of dissimilarities between the frames in the feature space were calculated using the Euclidean distance measure. 500 landmark points were obtained from farthest point sampling and subjected to minimization of the stress function. The switching distance was chosen equal to 2.5 for all *N-*glycans, which lies roughly in the middle of the range of distances characterized by Gaussian fluctuations (Figure S6). We set *A* = *B* = 4 in the high-dimensional space, and *a* = *b* = 2 in the low dimensional space, as defined by Ceriotti and co-workers^33^. The tolerance for the conjugate gradient minimization was set to 10^−3^, using 20 grid points in each direction and 200 grid points for interpolation. 5 annealing steps were used and the remaining trajectory data were projected on the constructed sketch-map.

### Kernel principal covariates regression model

The recently developed kPcovR algorithm was employed as described in the tutorials at https://github.com/lab-cosmo/kernel-tutorials, using the scikit-cosmo implementation. Besides the feature matrix **X**, the conformer strings assigned to each frame were employed as properties in the property matrix **Y**. Prior to fitting, the input was centered and standardized by removing the mean and scaling the data to obtain a unit variance. We note that our data set was used as a whole and not split into separate training and testing data sets. A Gaussian kernel with *γ* = 1 was used and mixing parameters *α* were calculated for each simulated glycan on a subset of 1000 frames for two dimensions, leading to values of 0.1 for M5, 0.5 for FM5, 0.9 for M9 and 0.1 for A2G2S2. The regularization parameter *λ* was set to 10^−4^ for the linear regression part. The error of the linear regression part was assessed by comparing the true properties **Y** and the predicted properties **Ŷ**.

### J-coupling calculations

Validation of the glycan structures obtained from MD simulations was carried out by comparing theoretically calculated with experimentally measured scalar ^3^*J*_*H,H*_ NMR coupling constants. The comparison is meaningful only for *ω* torsion angles in *α* 1 − 6 linkages, since *ϕ* and ψ lack the necessary proton pair, whereas the *J*_*H*5,*H*6_ and *J*_*H*5,*H*6′_ constants can be both computed and measured^60^ (see Fig. 7 below for the atom nomenclature). We note that caution must be taken when comparing the results of different force fields because of the inconsistencies in atom labelling conventions. In particular, the H6 (H6S) and H6’ (H6R) hydrogens are named ‘H61’ and ‘H62’ in CHARMM36m, respectively, while the opposite names (‘H62’ and ‘H61’) are used in GLYCAM06j. The theoretical calculations of the coupling constants was performed using three different sets of modified Karplus equations^61^, namely:

1. The equation of Altona and Haasnoot^62^:

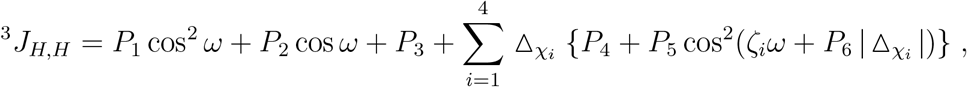

where the sum runs over the different substituents (in our case, H, C and two O), the *P* parameters are taken from the original data set, the electronegativity values 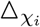 are equal to 0 for H, 0.4 for C and 1.3 for O and the substituent orientations *ζ*_*i*_ are either −1 or 1. The equation is applied to the torsion angles *ω* = H5–C5–C6–H6 or *ω* = H5–C5–C6–H6′. For example the former has electronegativity values of 0.4 for i=1, 1.3 for i=2 and 3, 0 for i=4, with *ζ*_1,2_ = 1 and *ζ*_3,4_ = −1.
2. The equations of Stenutz^63^:

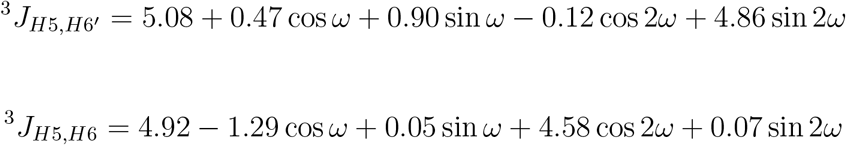

with *ω* = *O*5 − *C*5 − *C*6 − *O*6.
3. The equations of Tafazzoli^64^:

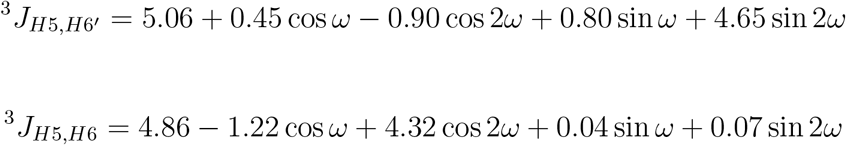

with *ω* = *O*5 − *C*5 − *C*6 − *O*6.

The general equation of Altona and Haasnoot can be adapted to different kind of linkages due to the flexible choice of substituents. In contrast, the equations of Stenutz and Tafazzoli were derived specifically from *J-*coupling constants computed with density functional theory for a model aldopyranosyl ring and D-glucose/D-galactose. The computed coupling constants were averaged over all 62500 frames in each REST-RECT simulation of the *N-*glycans (considering only the starting conformation s1). Block averaging was used to compute error bars, as described above for the probability distributions.

### Puckering

The general Cremer-Pople pucker coordinates *θ* and *ϕ* for 6-membered rings were used to evaluate the ring distortions (Figure 1, bottom), using the Puckering module of PLUMED (version 2.6). Two-dimensional histograms with 200 *×* 200 bins were constructed from the frames of REST-RECT simulations (only for the s1 starting conformation) and converted to free energy surfaces as described above. Two-dimensional pucker plots along *ϕ* and *θ* were plotted using the Mollweide projection, also termed homolographic or elliptical projection. This pseudocylindrical map projection is equal-area, meaning that areas, densities and, thus, free energy values are preserved.

## Results

In this section we first applied the REST-RECT method to different *N-*glycans, and benchmark its ergodicity against plain MD employing the CHARMM36m force field. In a second step, we compared the ability of different dimensionality reduction techniques, namely PCA, diffusion map, sketch-map and kPcovR, to effectively cluster distinct *N-*glycan conformations. The sampling and clustering protocol was then employed to compare the free-energy landscapes of different *N-*glycan systems, to assess the quality of different force fields validating the results against measured NMR J-coupling parameters, and to investigate the dependence of ring puckering on the glycan conformation.

### Comparison of REST-RECT and MD

Enhanced-sampling REST-RECT and plain MD simulations were performed to explore the conformational phase space spanned by the torsional angles of the *N*-glycan models shown in Figure 1 A. A fair comparison of the two methods was ensured by using the same total simulation time, amounting to 6 *µ*s for plain MD and to 500 ns for the REST-RECT simulations, which included 12 system replicas. The distribution of replica temperatures led to uniform and adequate exchange probabilities, as indicated by the achievement of at least 10 round-trips per replica in all cases (Figure S3). The obtained probability distributions of the conformer populations are shown in Figure 3 for the representative cases of the M5 and M9 glycans. Labelling of conformers, in line with the official IUPAC nomenclature for dihedral angles^56^, was performed as described in detail in the Method section (see also Figure S1). The first notable result is that stable and consistent conformer population distributions were obtained already after about 100 ns of REST-RECT simulation, irrespective of the starting conformation s1 or s2 (see Methods), as shown in the upper panels of Figure 3. In contrast, plain MD simulations displayed large fluctuations, poor convergence and significant dependency upon the starting conformation for individual conformers. This resulted in much larger error bars associated with the conformer distribution histograms (Figure 3 lower panel) in comparison with the enhanced-sampling simulations, especially in the case of the M9 glycan. Furthermore, plain MD simulations predicted different conformer populations than REST-RECT. For M5 the differences were not dramatic, and in particular the most-populated clusters corresponded to the same conformer. For M9, however, the ψ angle in the main 1–6 linkage between two mannose residues (gray box in Figure 3, lower panel) remained stuck in a *G*_ free-energy minimum and did not reach the global-minimum *T* conformation predicted by REST-RECT. Analogous conclusions were drawn for the other *N-*glycans FM5, A2G2S2, A2G2 (Figure S2), with FM5 showing the largest improvements associated with proper REST-RECT sampling.

The analysis revealed interesting common patterns of torsion-angle conformations in certain structural elements across the investigated models. For instance, the sequence *G*_−_*A*_+_*G*_−_*A*_+_ was predicted as the global minimum of the chitobiose core for all glycans, and the *G*_+_*Tgg* sequence characterizes the 1–6 linkages in most cases. Moreover, there was an evident preference for a *gg* conformation of the *ω* angle, which originates from the *gauche* effect.

### Dimensionality reduction of *N-*glycans

The set of strings associated with the most-populated conformer clusters of a given *N-* glycan (as shown in Figure 2) is a reduced representation of the free-energy minima in the high-dimensional conformational phase space spanned by all torsion angles. However, such representation becomes cumbersome when comparing different *N-*glycan systems, and does not give a measure of the structural differences among the different conformers. We therefore investigated the ability of several dimensionality reduction techniques to deliver two-dimensional representations of the conformer clusters in an efficient and physical meaningful manner, using all torsion angles as input features. As stated in the Introduction, we concentrated on three unsupervised-learning algorithms, namely PCA, diffusion maps and sketch maps, and the supervised-learning algorithm kPcovR.

PCA and diffusion map generated almost identical two-dimensional representations (Figures 4 and S7) and gave similar eigenvalue progressions along the PCA/Diffusion components (Figure S6). For all *N-*glycans, there was an obvious gap between the first one to three eigenvalues (two for the case of M5 shown in Figure 4) and the remaining ones, indicating that a few corresponding structural features are more important than others in differentiating the glycan conformers. Having a closer look at the PCA of M5, *PC*1 differentiates conformers along their *ω* torsion angle in the side branch 6–, and *PC*2 along the main branch **6**–, corresponding to 25.0 % and 19.5 % of variance in the underlying data, respectively. Knowing that the highest variance is included in the rotations around *ω* torsion angles gave us an a-posteriori justification for the two selected initialization states s1 and s2, differing in the states of *ω* in **6**–, situated in very different energetically conformers. This differentiation of conformers from each other by two *ω* angles gave rise to four main groups of conformer clusters (Figure 4). The overlap between the conformer clusters in each of those groups originated from the fact that these conformers only differ in ψ angles, which can not be resolved in this two-dimensional representation, revealing the limitations of the PCA and Diffusion map algorithms (Figure 3).

**Figure 4:**
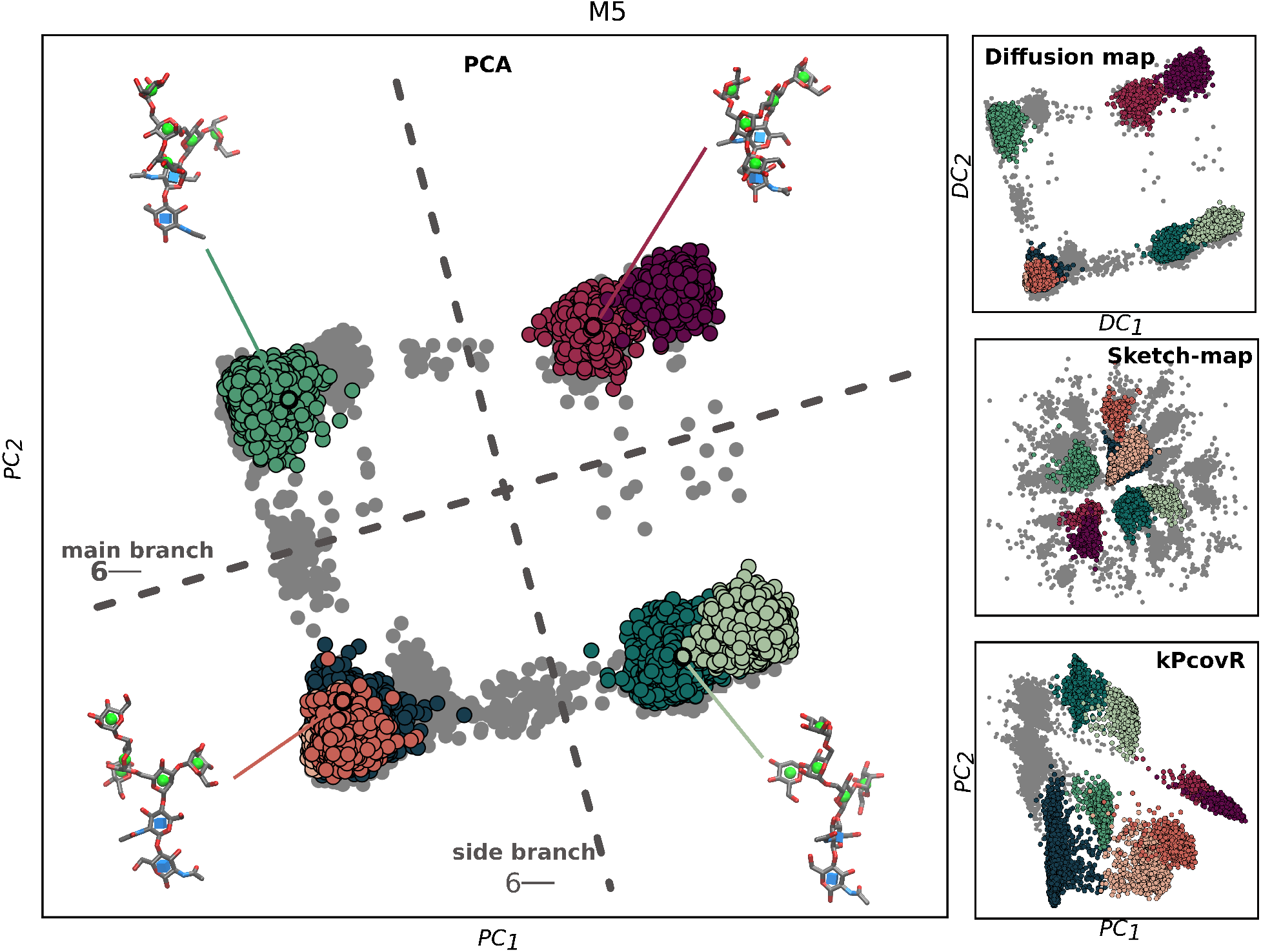
Comparison of four different dimensionality reduction algorithms to cluster the distinct conformers of *N-*glycan M5. Principle component analysis (PCA), diffusion map and sketch-map employ all M5 torsion angles as features, whereas kernel principle covariates regression (kPcovR) additionally uses the conformer strings as additional property. Each gray point corresponds to one frame and colored points to the respective conformers given in Figure 3. Sampling was performed from REST-RECT simulations (only s1).

In fact, it is interesting to note that the number of highest, well-separated PCA or Diffusion-map eigenvalues is equal to the number of *ω* angles present in the *N-*glycan structures for the M5 (two), FM5 (three) and A2G2S2 (one) glycans (Figure S6). In this respect, M9 is an exception, presenting only one well-separated eigenvalue (corresponding to the 6– branch), but two *ω* angles. Correspondingly, in the two-dimensional maps there is a clear separation of cluster conformers along the *PC*1 component, but some overlap along the *PC*2 component (Figure S7).

The Sketch-map algorithm clustered conformers in a similar way to PCA and diffusion map, differing in the overall spatial arrangement, but with only marginally better separation of the clusters. The Sketch-map analysis did not allow for a ranking of the components, and thus for an unbiased identification of the most important structural features of the system. The kPcovR algorithm separated the most-occupied conformers in the most effective way, resulting in cluster clouds with only little overlap to neighboring ones (Figure 4). However, the algorithm did not allow for a meaningful interpretation of how conformers are separated or clustered together, since no characteristic feature could be assigned to the kPcovR principal components. It rather seems that the clusters were ranked according to their population probabilities along *PC*1, as suggested by the progression of colors from left to right in the two-dimensional map and in the histograms of conformer probabilities (Figure 3 and Figure 4). The calculated losses for the regressions in kPcovR can only be interpreted as relative values, as **Y** consists of discrete conformers, but the linearity between **Y** and **Ŷ** along the target (red line) is very clear (Figure S6).

From this analysis, we conclude that, for the investigated systems, PCA, diffusion map and sketch-map can be used almost interchangeably with respect to their physical meaning, while kPcovR may be useful whenever a two-dimensional representation with well-separated conformer clusters is sought for. In this study, we employed PCA in the following applications of the method, as it is the most straightforward and computationally effective algorithm.

### Comparison of *N-*glycan structures

As a first application, we compared the conformational landscape of high-mannose-type and complex *N-*glycans. To this purpose, we used PCA as a method to represent the free-energy maps associated with the conformational ensembles of our five glycan models in two dimensions. We focused on the influence that various chemical modifications (fucosylation, sialylation, variation of branch length) might have on the resulting structures. In doing so, we included in the analysis only the structural features common to all compared structures, highlighting the effect of the mentioned modifications on the free-energy maps (Figure 5). In general, the analysis showed that the core structure of the high-mannose-type glycans was only marginally affected by addition of further residues. Fucolsylation of M5 (FM5) had no effect at all on the conformational free-energy landscape, whereas elongation of the shorter branch by two units (M9) led to a slight stabilization of the main conformer and destabilization of the secondary minima at larger *PC*1 values (Figure 5, upper row). Instead, sialylation of A2G2 (A2G2S2) with additional Neu5Ac units on both branches deepened slightly all three secondary energy minima at the expense of the most-populated region of the conformational phase space in the bottom-right corner of the map (Figure 5, lower row).

**Figure 5:**
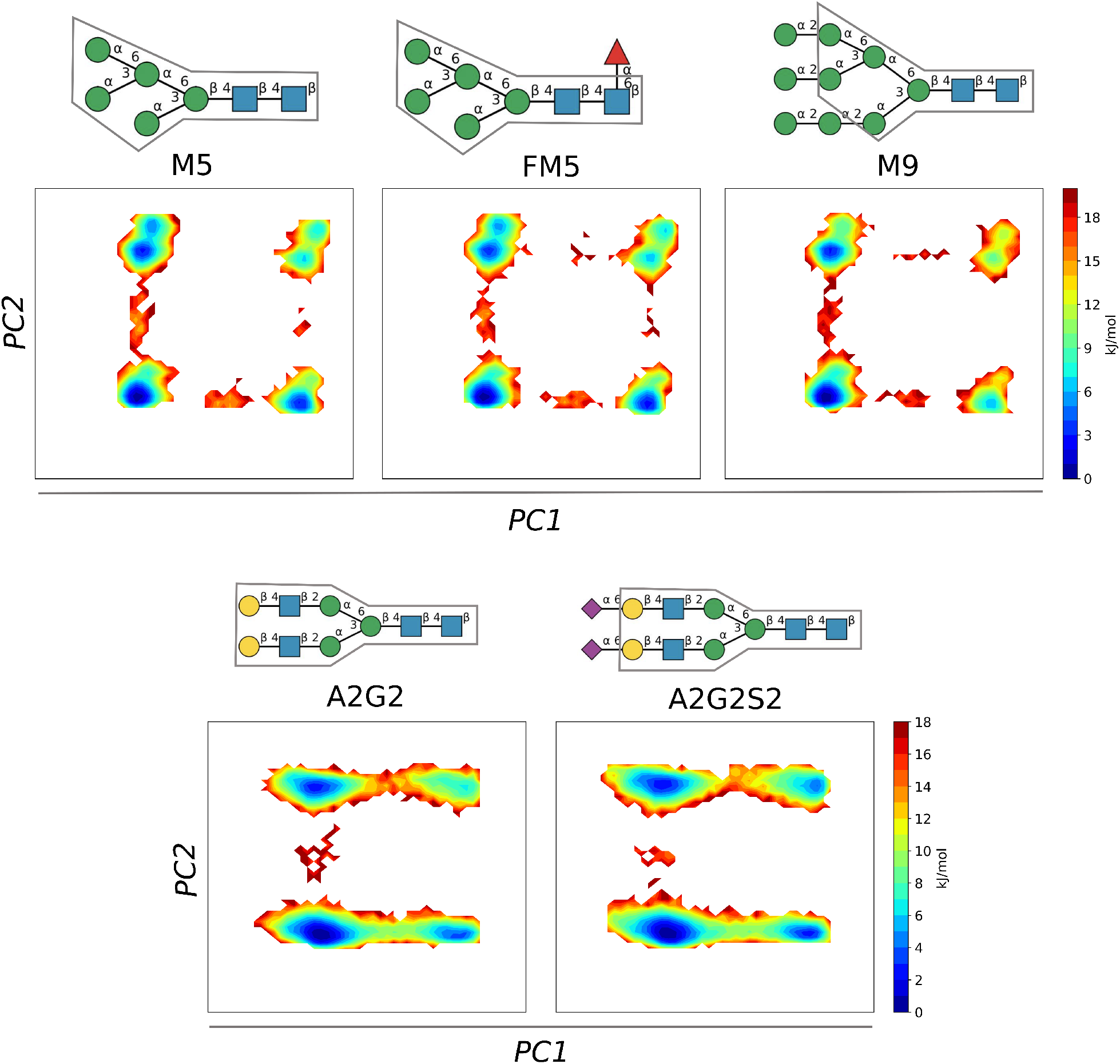
PCA free-energy maps of different *N-*glycans. The upper panels compare the three high mannose type *N-*glycans M5, FM5 and M9, whereas the lower panels compare A2G2 against its sialylated variant A2G2S2. Only the torsion angles common to all structures (boxes around the schematic glycan models) where used as features in the analysis. A common PCA was performed by concatenating the datasets of M5, FM5, M9 and those of A2G2 and A2G2S2, respectively. Sampling of the phase space was performed with REST-RECT simulations, starting from the s1 conformation.

### Comparison of force field predictions

Besides enabling comparisons of different *N-*glycans structures, the developed methodology allowed us to compare very accurately the structural prediction capability of different force fields. Here, we compared the two most widely used force fields for protein and carbohydrate systems, namely CHARMM36m and GLYCAM06j. The results for A2G2S2 are shown exemplarily in Figure 6, and for the other glycans (except FM5 because of its similarity to M5) in Figure S8. The comparison revealed very substantial differences in terms of both conformer distributions and free-energy landscapes, and even the global-minimum structures were different. CHARMM36m predicted that the majority of conformers (and thus the global free-energy minimum) cluster on the right-bottom region of the PCA map, in contrast with GLYCAM06j, which predicted a global minimum in the left-bottom region. The position of the secondary minima was also different in the two cases. Comparison of the predicted conformer strings indicated that the major differences arise from the ψ angle of the main branch **6**– as well as the two *ω* angles of the terminal 1-6 linkages between Gal and Neu5Ac (Figure 6, lower panels). In the global-minimum structure predicted by CHARMM36m ψ was in a *T* conformation and *ω* in a *gt* conformation. These conformations changed to *G*_+ and *tg* in the global-minimum structure predicted by GLYCAM06j, respectively. Similar considerations hold for the M5, M9 and A2G2 (Figure S8), although the conformer distributions were less dramatically different than in the case of AG2S2.

**Figure 6:**
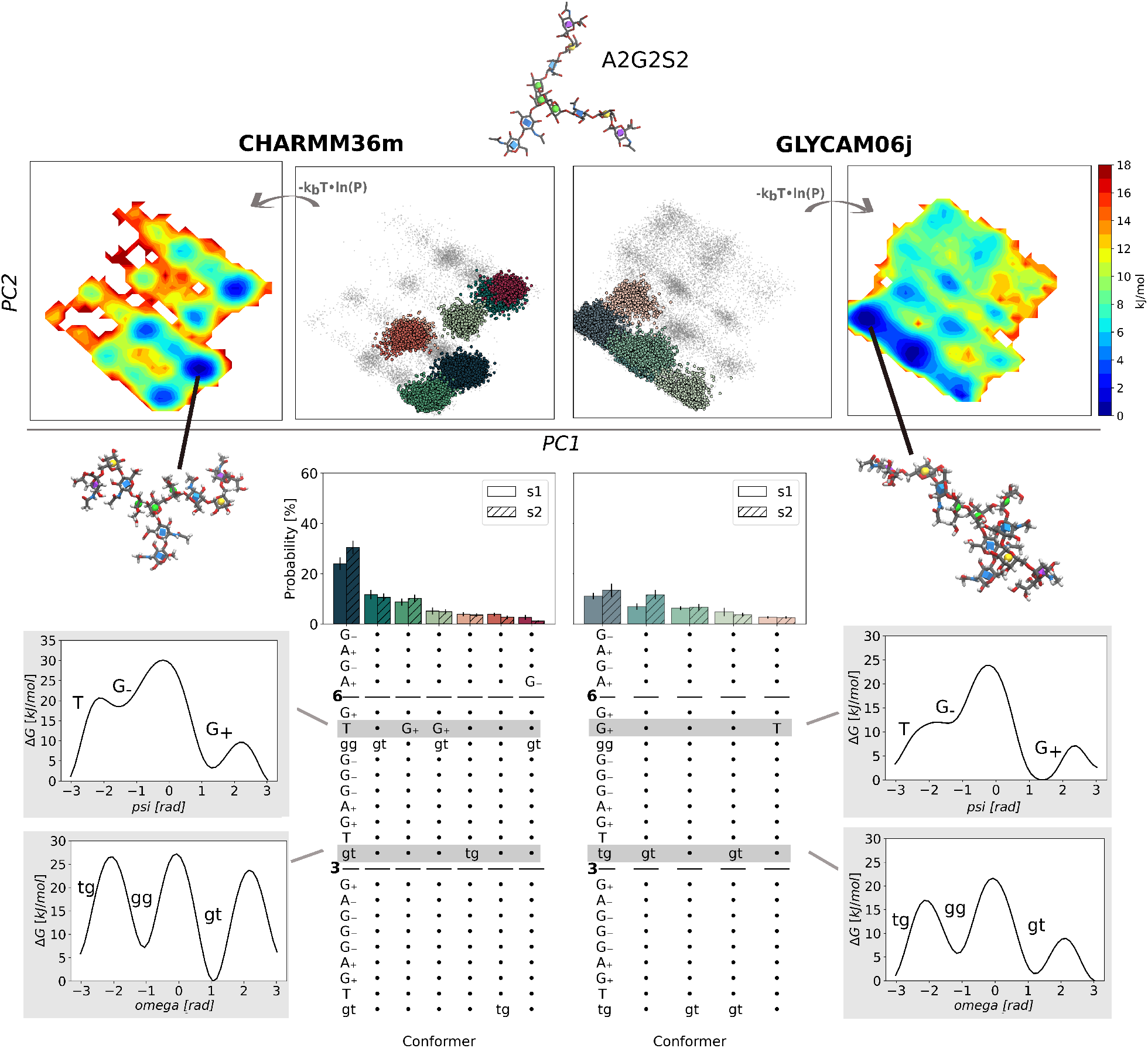
Comparison of the conformational ensembles of A2G2S2 predicted by REST-RECT simulations with either the CHARMM36m or the GLYCAM06j force field. The upper panels show the PCA maps of conformer clusters and the corresponding free-energy landscape. The clusters are colored in accordance to the conformer distributions shown in the lower panels. Free energy profiles along selected torsion angles (indicated by the gray rectangles) are represented besides the conformer strings and labeled with the conformations of the free energy minima.

By looking at the one-dimensional profiles along selected torsion angles, it becomes evident that the discrepancies arise from only subtle differences in the force field parametrizations. For instance, the free-energy differences between the *T* and *G*_+ conformations of the ψ angle, or between the *gt* and *tg* conformations of the *ω* angle, were less than 5 kJ/mol. However, such small differences have a profound effect on the resulting multi-dimensional free-energy landscape, and lead to rather distant global minima, as observed above. In the next section, we will show that these differences led to markedly different predictions of NMR spectroscopic fingerprints of the glycan populations by the different force fields.

### Force field validation by calculation of NMR parameters

A comparison of experimental NMR data with the corresponding observables predicted theoretically by the two force fields was performed to ascertain which one can be considered more accurate in terms of torsion angle description. We stress that this assessment is only reliable when the ergodicity of the simulations is fulfilled, which requires complete phase-space sampling by means of converged REST-RECT simulations.

As shown above, the largest variability among the different conformers of *N-*glycans originated from the *ω* torsion angles around the 1–6 O-glycosidic linkages. The three protons H5, H6, and H6’ harbored by these linkages (see Figure 7) gave rise to well-defined NMR J-coupling constants, whose values depended on the relative distances between the H nuclear spins, and thus on the conformation of the *ω* angles. Therefore, a comparison between measured and theoretically computed *J*_*H*5,*H*6_ and *J*_*H*5,*H*6′_ frequencies enabled a clear validation of the predicted glycan structures^60^. The calculations were performed by ensemble averages of the coupling constants computed for all conformers sampled by the REST-RECT simulations, using three different parametrizations of the empirical Karplus equation, as described in the Methods section (Figure 7).

**Figure 7:**
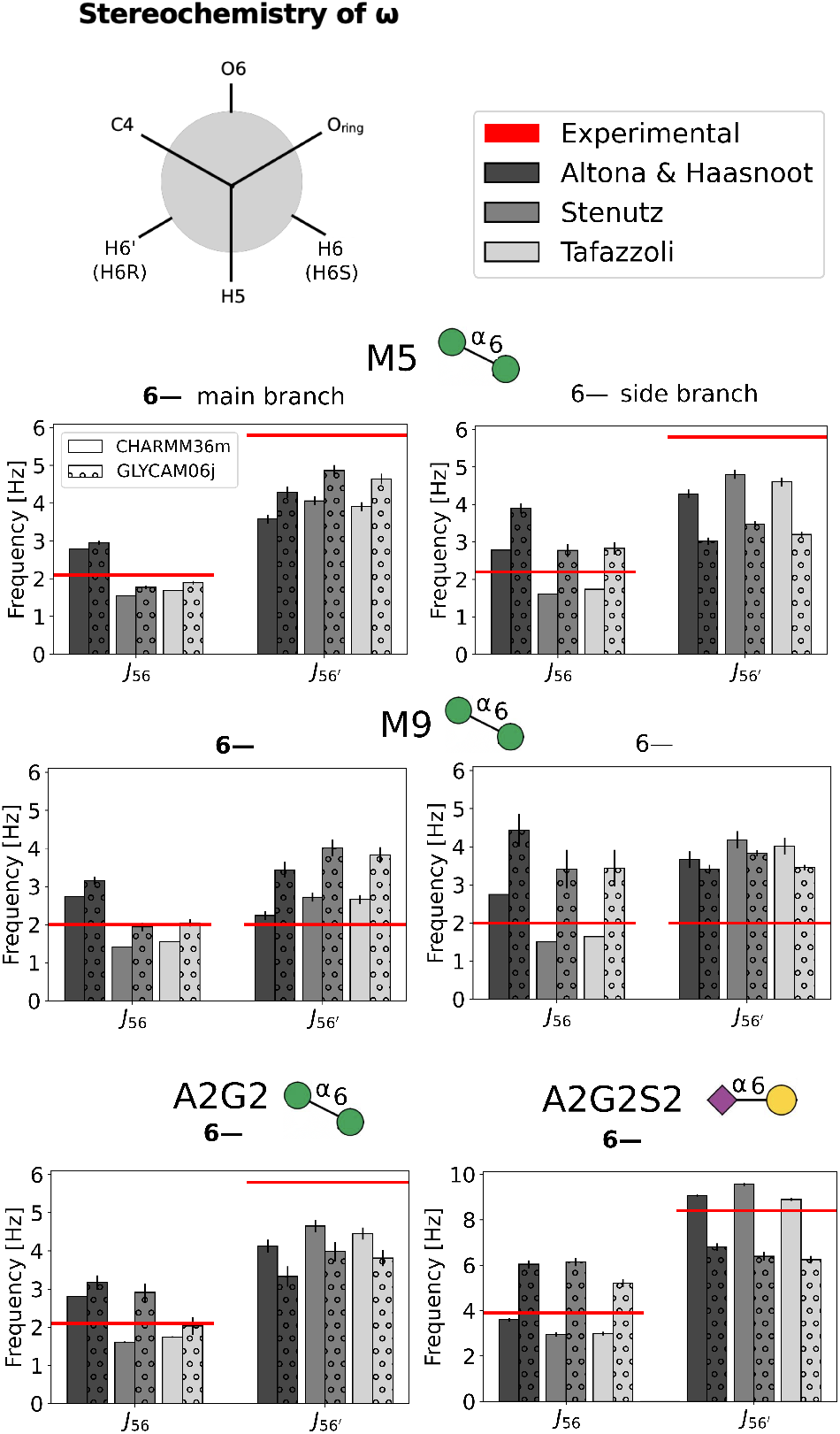
Validation of the *ω* angle populations by comparison of computed and experimental NMR J-coupling constants for M5, M9, A2G2^60^ and A2G2S2^27,66^. The upper-left panel shows the stereochemistry of an *ω* angle in a *gg* conformation along its C5 and C6 atoms, with labeled protons. The legend on the right reports the color code of the plots below, referring to the three different parametrizations of the Karplus equation used to compute the J-coupling constants. All plotted values are also reported in Table S1.

For the main branch of M5, GLYCAM06j led to better agreement between experimental and theoretical *J*_*H*5,*H*6_ and *J*_*H*5,*H*6′_ frequencies, whereas CHARMM36m performed better for the side branch. Regarding M9, CHARMM36m performed better than GLYCAM06j for both *ω* angles, although both force fields overestimated the *J*_*H*5,*H*6′_ frequency by over 2 Hz. We would like to note that the experimental J-couplings of M9 were only reported as approximations in the original paper^60^, but used here due to the lack of other data sources. For the only *ω* torsion angle in A2G2, CHARMM36m predicted slightly better frequencies than GLYCAM06j. For the sialylated variant A2G2S2, no experimental parameter of the *ω* torsion angle between two Man residues was available, therefore the comparison was made for the J-coupling constants of the 1–6 linkage between Gal and Neu5Ac. In this case, the experimental values were collected for a different glycan, namely trisaccharide sialyl-*α*-(2-6)-lactose,^27,66^ but could be used here as an approximation, because this glycan carries the same terminal branches as A2G2S2. The predicted CHARMM36m values of both *J*_*H*5,*H*6_ and *J*_*H*5,*H*6′_ were in very good agreement with the experimental ones.

Overall, neither force field reproduced all experimental J-coupling constants with great accuracy (i.e., within the intrinsic error bars of the theoretical method), but CHARMM36m seemed to deliver better structural predictions than GLYCAM06j, especially in the cases of M9 and A2G2S2.

### Analysis of the sugar puckering conformations

So far we only focused on the torsion angles of the *N-*glycan molecules, which were considered as explicit RECT collective variables in our simulations. However, the combination with the REST2 method allowed also good sampling of other structural degrees of freedom, in particular of the puckering conformations of the individual monosaccharide units. In this section we investigate whether the puckering free-energy landscapes spanned by the Cremer-Pople parameters *θ* and *ϕ* were dependent on the specific conformation of the glycan, as determined by its torsion angle.

To this aim, we constructed and represented two-dimensional polar free-energy maps in a way that conserves the area defined by intervals of the *θ* and *ϕ* pucker coordinates. As usual, the free energy was computed from the histograms of conformer population probability in the ground replica of the REST-RECT simulations. In Figure 8 we show the puckering maps of the GlcNAc1 and GlcNAc2 units of the most-populated and the second-most-populated torsion angle conformer clusters (with respect to the distribution reported in Figures 3 and S2).

**Figure 8:**
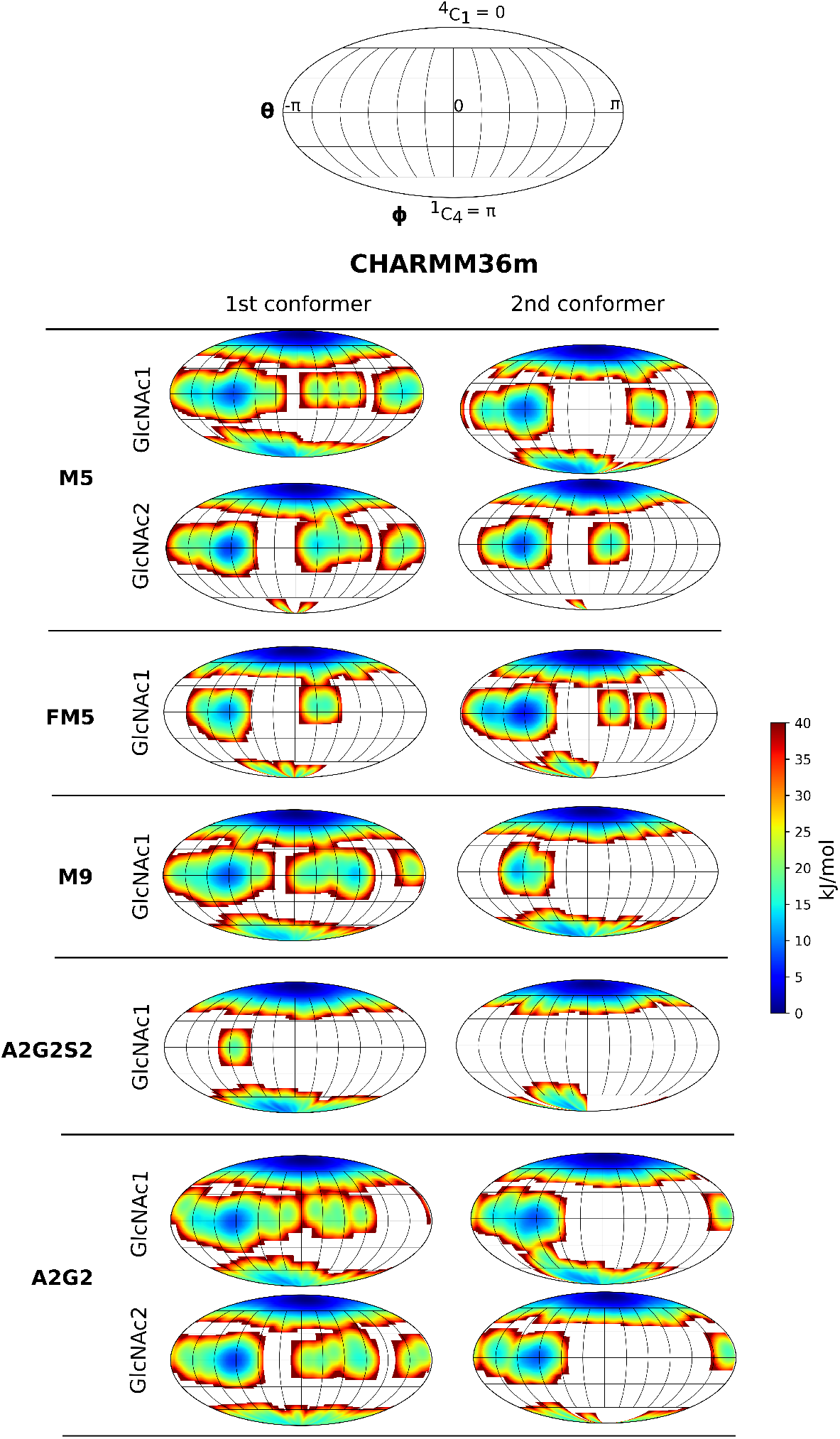
Free energy surface along the Cremer-Pople puckering coordinates *θ* and *ϕ* for GlcNAc residues of all *N-*glycans compared for their most and second most occuring con-formers. Collective variables were computed from REST-RECT simulations by histogram construction and convertion to free energies.

For all considered glycans, the CHARMM36m force field predicted a ^4^*C*_1_ chair conformation as the global minimum of the monosaccharide units in both conformer states. Minor differences could be observed in the relative position (or even appearance) of the secondary minima corresponding to boat conformations (at the equator of the maps, for *θ* = *π/*2), and mostly for monosaccharide units situated in the core of the *N-*glycans. Third-most and fourth-most conformers have also been analyzed, but not explicitly shown as they resemble the same occupaction of minima as seen in the most-populated and the second-most-populated clusters. Whether larger shifts of the local minima or even stabilization of boat conformations could result from constraining the glycans within conformer states far from their natural global minima (e.g. upon binding of the molecule into tight protein pockets) is a question that should be investigated in future works.

Analysis of the pucker landscapes along the branches of the A2G2S2 glycan, which is composed by very diverse monosaccharide units, revealed again a strong propensity for the chair conformation ^4^*C*_1_, except for the terminal Neu5Ac units that were in a ^1^*C*_4_ conformation (Figure S10). However, the relative boat propensities were quite different for the different units, being very low or absent for Man and Gal units, more evident for GlcNAc units and strongest for Neu5Ac units. We note that the GLYCAM06j, in comparison with CHARMM36m, generally predicted a broader exploration of the pucker phase-space, resulting in an increased appearance of local minima and smaller energy differences between different regions of the maps (Figure S10). Only the terminal Neu5Ac units presented very similar maps for both force fields, with the same distribution of minima and the same degree of phase-space exploration.

## Discussion

Glycosylation is an ubiquitous modification of biomolecular systems, involving proteins, lipids and, as more recently discovered, RNA^67^. As a prominent example, glycans tethered to the SARS-CoV-2 spike protein modulate its enzymatic function^68,69^, providing but a small glimpse of the importance of glycosylation in life science^3^. However, the tremendous diversity of *N-*glycan structures promoted by all possible cellular glycosylation pathways hampers a rational understanding of clear structure-function relationships at the basis of a still misterious sugar code^70^. Our claim is that a correct and comprehensive description of the whole ensemble of conformers is crucial for the prediction of the biological function of glycan systems, motivating a fundamental exploration of the conformational phase space of representative *N-*glycans by means of enhanced-sampling molecular dynamics simulations (Figure 2).

We were able to show that REST-RECT simulations provide converged conformer distributions with complete sampling of all torsion and puckering angles within a few hundred nanoseconds of cumulative time, with better accuracy and using less computational time than long plain MD simulations. Sufficiently short round trip times reveal the strength of the RECT method, capable of biasing up to 22 CVs simultaneously while still ensuring adequate diffusion in the replica space. This behaviour originates from an adjusted ergodicity by scaled bias factors over the replica ladder, ensuring a proper compensation of free energy barriers by a self-consistent addition of one-dimensional bias potentials^28^. Alternative methods such as temperature REMD^19^ are computationally too demanding in solvated systems ^71^. Bias-exchange metadynamics^72^ would have required more replicas for the same number of biased CVs, and would have only enabled biasing them one at a time^28^. It may be argued that some of the chosen CVs are in fact redundant; in particular, axial *ϕ* torsion angles in *α*-linkages occupy only the gauche conformation (G_+_) due to the so-called exoanomeric effect. This is due to the favorable overlap of one oxygen lone-electron pair with the antibonding orbital of the adjacent C–O bond ^14,73,74^, an effect that needs to be appropriately mapped by torsion, Lennard-Jones and Coulomb terms in force-field potentials. However, there is no computational advantage in excluding those CVs from the biased scheme, and is indeed reassuring to see that the results do confirm such background-knowledge details.

Whereas the computational method ensures ergodicity of the performed simulations, their accuracy in predicting experimental observables remain limited by the functional form and parametrization of the employed force fields. Various carbohydrate force fields have been developed and refined for selected systems and cases, such as the stability of protein-carbohydrate complexes, the conformational behaviour of linear polysaccharides, or the ring distortions of monosaccharide units^15,16,75^. Generally speaking, the main families of biomolecular force fields, namely CHARMM, GLYCAM (AMBER) and GROMOS, have been shown to have good performance in reproducing the behavior and predicting experimental data of polysaccharide systems, with few exceptions^14,16^. However, depending on the saccharide size, the property under investigation and the required levels of detail, differences among the force field families do emerge, which can be traced back to how well the steric, electrostatic, and torsional energy terms represent the physical reality and mimic the actual glycan behaviour^14^.

We have performed an in-depth analysis of the *N-*glycan conformer distributions predicted by the CHARMM36m and GLYCAM06j force fields, focusing on converged free energy profiles of torsion angles that shape the three dimensional glycan structure and its flexibility. The observed different phase-space distributions for A2G2S2 and A2G2, as well as the different conformer distributions for M5 and M9, originate from different free energy profiles around the ψ and *ω* torsion angles in 1–6 linkages. Especially the ψ angle of branch **6–** in A2G2 and A2G2S2 is a critical feature, which differentiates between two main conformers, previously named ‘backfold’ and ‘extended’^26^. CHARMM36m consistently produced conformer distributions with only a few high populated states, whereas GLYCAM06j produced broader distributions and flatter associated free-energy landscapes. The frequent revisions of the force-field parametrization of torsional terms, up to the present days, indeed shows that a correct description of rotational barriers in glycan systems is not at all straightforward^76^. In particular, the contribution of torsional energy and electrostatic interactions needs to be balanced with great care^76^. The two force fields compared here differ especially in the latter term: CHARMM36m adjusts partial atomic charges to fit solute-water interactions of carbohydrate fragments computed with quantum mechanical methods, whereas the partial charges of GLYCAM06s are derived from the restrained electrostatic potential (RESP) method^76^.

Our comparison of computed J-coupling constant values to the sparsely available data from NMR experiments indicated an overall better performance of the CHARMM36m force field. The three tested parametrizations of the Karplus equation yielded consistent results, although they are all based on empirical parameters so that some discrepancies are both expected and unavoidable. While all of our calculations were performed with the TIP3P water model, more complex water models like TIP5P have been shown to impact the predicted carbohydrate aggregation and protein-carbohydrate interactions^77–79^. Therefore, future studies should also focus on a critical assessment of how different solvent models impact the performance of glycan force fields in predicting the correct conformer cluster distributions.

The here-examined *N-*glycans have been the subject of several previous investigations. Both experimental and computational studies of M9 suggested that it is mainly confined in a gauche conformer, meaning that the *ω* torsion angle in the **6–** branch should be in a *gg* conformation, which is in agreement with our findings for both employed force fields^80–82^. The observed stabilization of the global minimum of M9 after elongation of one M5 branch by two 1–2-linked mannose units (see Figure 5) is probably due to an increased number of inter-branch hydrogen bonds^83^. A2G2S2 has been studied by Yang and coworkers^24^ using REST2 in combination with Hamiltonian bias potentials, employing the CHARMM36 force field. This approach is similar to REST-RECT, the only difference being that Yang and coworkers used a biasing profiles on the torsional angles as obtained in preliminary umbrella sampling simulations, whereas in our RECT scheme the compensating profiles are computed on the fly. In fact, the reported free energy profiles for individual torsion angles on ref.^24^ are overall in agreement with our CHARMM36m simulations (Figure S9), demonstrating converged phase-space sampling in both studies.

A2G2 was previously investigated by REMD using the GLYCAM06g force field, and the predicted relative populations of the *ω* angle (O6–C6–C5–C4) in branch **6–** amounted to 71 % and 28 % for the *gg* and *gt* conformers, respectively^26^. Our simulations with the GLYCAM06j force field, however, gave average values of 80 % for *gg*, 11 % for *gt* and 9 % for *tg*. While the force field versions g and j only differ in the atom labelling for consistency with other AMBER force fields or in the addition of parameters for protein-carbohydrate linkage, the simulations by Nishima and coworkers^26^ differ from ours with respect to the type of sampling method. We believe that our REST-RECT simulations provide a more complete phase-space sampling, as demonstrated by the very good convergence (Figure S4) and the clear independency from the chosen initial configurations. Other earlier investigations of A2G2 using the CHARMM36 force field revealed a distribution of 52 % *gg* vs 48 % *gt* conformations in the *ω* torsion angle (O6–C6–C5–O5)^18^. Our values of 71 % for *gg* and 29 % for *gt* computed with CHARMM36m, however, are closer to the experimentally estimated values of 65 % and 35 %, respectively^18,84^. As identical force field parameters were used in both studies, the associated differences can have multiple reasons, namely (i) the use of the sTIP3P water model in contrast to the mTIP3P model used here, again poiting towards the need of better assessing the performance of different solvent models; (ii) incomplete phase-space sampling in the earlier simulations; (iii) the fact that in their simulations Galvelis and coworkers prevented ring inversion for all monosaccharide rings, although there are hints about a possible influence of puckering states on the glycan linkage conformations ^85,86^.

The last point opens a further argument worth discussing. Ring flipping (inversion) or extended puckering distortion is a process that happens on the *µ*s time scale and therefore requires very long simulation times or enhanced sampling^14^. It was long considered irrelevant for most monosaccharides in polysaccharide chains, and therefore the puckering degrees of freedom were often ignored, reducing the complexity of MD simulations^18^. However, as mentioned above, several computational studies revealed the importance of ring flipping events in determining the polysaccharide conformer distribution^12,75,86^. For instance, GlcNAc and Neu5Ac were shown to readily undergo ring flips^87–89^, which could also be observed in our *N-*glycan systems, in contrast to the more rigid mannose residues (Figures 8 and S10). In particular, the puckering of saccharide units connected via 1–4 linkages within glycan chains was suggested to influence the *ϕ* and ψ distributions of the adjacent linkages.

While we do observe that puckering free energy landscapes of GlcNAc residues located in the chitobiose core differ among different conformers, we cannot confirm a direct effect on the adjacent linkages of puckering-prone units. In line with our observations, previous studies of glycan monomers and trimers did notice differences in the puckering landscapes predicted by different force fields, and in particular pointed towards a better performance of the CHARMM36m force field^75^. Accurate prediction of ring-inversion free-energies is expected to be very important for strongly constrained systems, such as glycan chains bound in protein pockets and subjected to enzymatic reactions, where ring-inversion is often a key step of substrate activation before e.g. hydrolysis of the adjacent linkages^90^.

Importantly, we report a systematic comparison of dimensionality-reduction techniques applied to *N-*glycans. Up to now, this has been limited to PCA using atom coordinates as input features^82,91^. Nevertheless, clustering of glycan conformers has been already performed in the past, using e.g. end-to-end distances between glycan branches to describe their flexibility and identify the main conformers^23^. Additionally, the usage of spherical coordinates was introduced to describe the dynamical behaviour of the **6–** branch, albeit the procedure was applied to the description of only one single branch^26^. Our successful application of various dimensionality-reduction techniques to represent the high-dimensional phase space of *N-*glycans highlights their enormous potential in delivering a consistent analysis of the most-populated conformer clusters, while simultaneously providing meaningful information about the most important structural features behind the used descriptive variables. This becomes very important for glycans composed of diverse monosaccharide units arranged in complex branched chains, presenting additional chemical modifications (fucolsylation, sialylation) and including more than two *ω* torsional angles, making structural relationships not as intuitive as for small glycans like M5.

Complementing the clustering and dimensionality reduction analysis, the here-introduced conformer strings provide a solid and IUPAC-nomenclature-compliant way of labeling different conformers. Comparison of the strings easily reveals differences and similarities, which are immediately traceable to specific linkages and torsion angles. Previous classifications of identified conformers were performed with less clear nomenclature rules. For instance, the groups ‘backfold’, ‘half backfold’, ‘tight backfold’, ‘extended-a’ and ‘extended-b’ were defined according to their ψ and *ω* torsion values of the first 1–6 linkage. These groups can still be differentiated using a low dimensional representation constructed by PCA (see e.g. Figure S11 for A2G2). In line with a previous study, we found that ‘half backfold’ conformers is in fact part of the ‘backfold’ cluster^18^. However, this classification system is again limited to the description of only one single branch.

Finally, we note that our developed analysis framework is not limited to the investigation of *N-*glycans, but can be extended to other glycan classes with other linkage types, such as O-glycans, GPI-anchors, or glycosphingolipids. Furthermore, the analysis can be extended to glycans bound to protein systems, although care must be taken in ensuring complete phase-space exploration despite the constraints dictated by amino-acid/glycan interactions. Adaptation of the REST-RECT algorithm to the study of protein-bound glycans will be the aim of future investigations.

## Associated content

The generation of conformer strings and plotting of conformer string distribution plots was done by GlyCONFORMER, a python-based package deposited on GitHub under https://github.com/IsabellGrothaus/GlyCONFORMER. An example dataset is given, demonstrating the analysis workflow in a jupyter-notebook. Additionally, the repository includes exemplary jupyter-notebooks offering help in performing dimensionality reduction algorithms. Employed PLUMED input files for REST-RECT simulations were deposited in the PLUMED-NEST repository^92^ under PlumID:22.028. Structure and trajectory files for all *N-*glycans simulated with the two employed force fields, CHARMM36m and GLYCAM06j, can be accessed from https://doi.org/10.5281/zenodo.6542267.

## Acknowledgement

The authors thank Georg Bossenz for the support implementing a python script to calculate J-coupling values from MD simulations. Computational resources were provided by the North German Supercomputing Alliance (HLRN), project hbb00001. Additionally, computational resources were provided by the supercomputing centre CINECA, in the framework of the HPC-Europa3 Transnational Access programme under application HPC177SW8P.

## Supporting Information Available

The Supporting Information is available free of charge in a separate pdf file. It includes supplementary figures and tables referenced in the main text.

