## Supplementary Information for "Exploration, representation and rationalization of the conformational phase-space of N-glycans"

#### Contents

Conformer string . . . . . . . . . . . . . . . . . . . . . . . . . . . . . . . . S-2

Comparison of REST-RECT and MD . . . . . . . . . . . . . . . . . . . . . . . S-3

CHARMM36m . . . . . . . . . . . . . . . . . . . . . . . . . . . . . . . . S-3

GLYCAM06j . . . . . . . . . . . . . . . . . . . . . . . . . . . . . . . . S-8

Dimensionality reduction of N-glycans . . . . . . . . . . . . . . . . . . . . . S-12

Force field comparison . . . . . . . . . . . . . . . . . . . . . . . . . . . . S-14

Free energy profiles . . . . . . . . . . . . . . . . . . . . . . . . . . . . . S-17

Force field validation via NMR parameter . . . . . . . . . . . . . . . . . . . S-19

#### Conformer string

Figure S1: Conformer string construction based on torsion angle configurations for various $N$-glycans. Rules for the assignmemt of a conformer string are given and two examples shown.

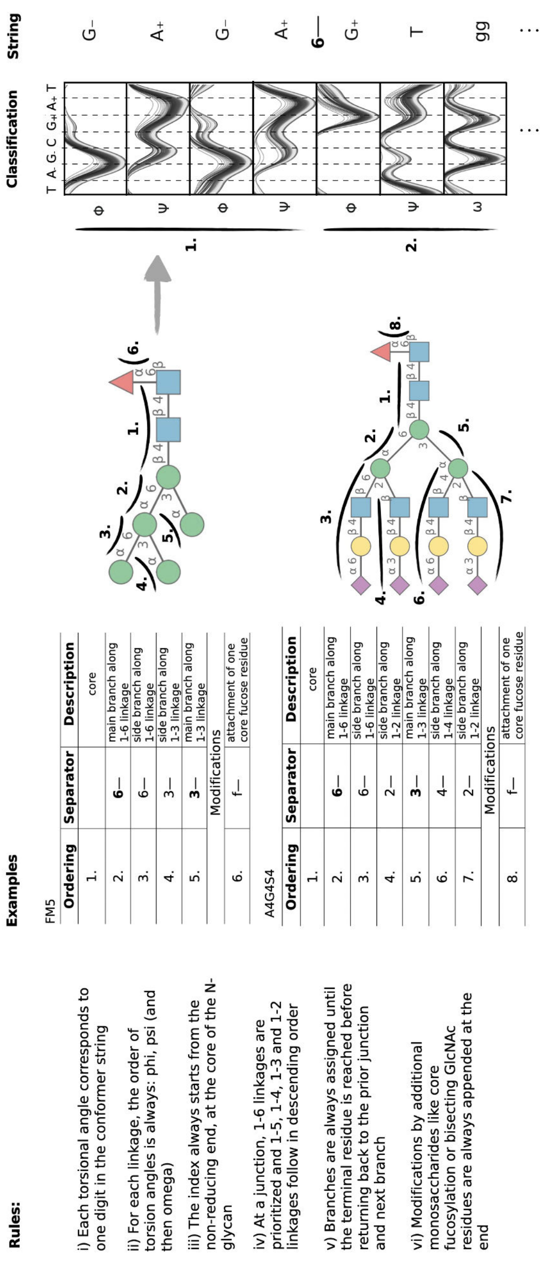

### Comparison of REST-RECT and MD

#### CHARMM36m

Figure S2: Comparison of REST-RECT with plain MD simulations for systems FM5, A2G2S2 (and A2G2) employing the CHARMM36m force field (FF). The upper panel includes atomistic structures of each glycan. The middle panel shows the moving average for the three most populated conformers using a window size of 100 ns (REST-RECT) and 1.2 $\mu$s (MD), corresponding to the same sampling time. Two separate simulations were performed with differing initial starting configurations (s1,s2). The lower panel represents conformer distributions for both starting structures. The conformer string is given on the x-axis, where each digit stands for a torsion angle, the letter representing the occupied minima. Dots are used instead of letters when no change in occupancy could be observed in comparison with the most occupied conformer.

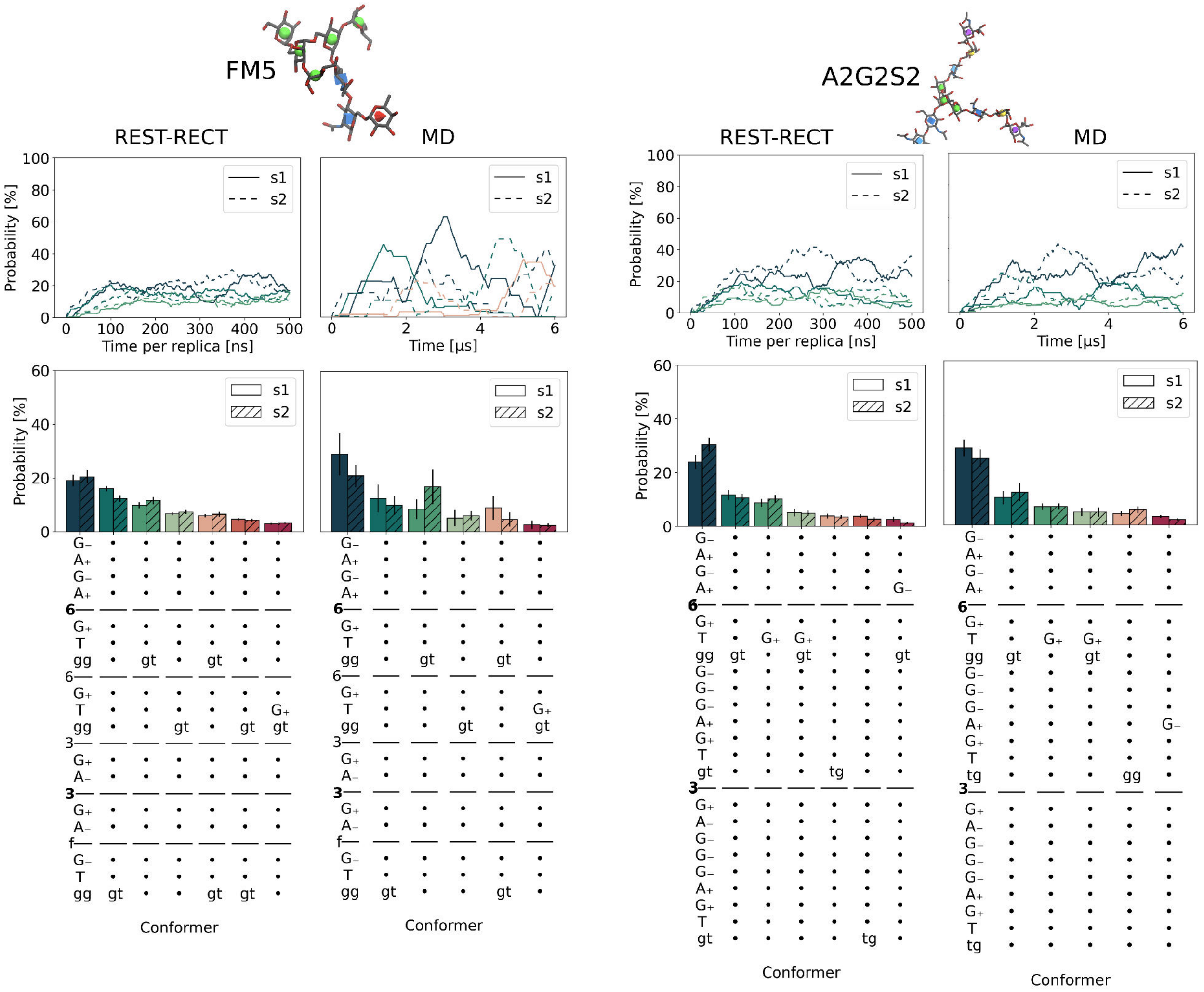

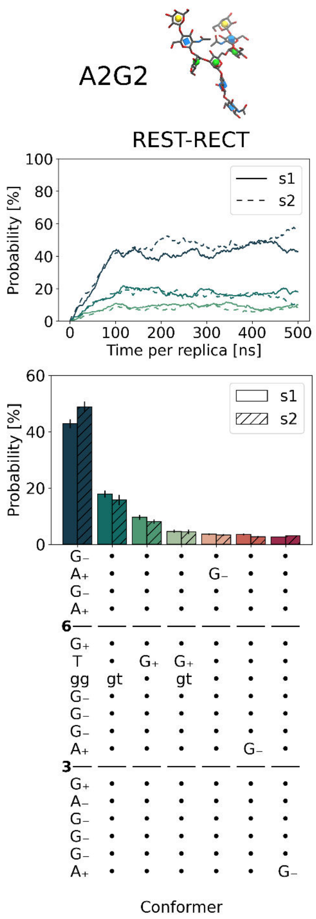

A2G2
REST-RECT
Probability [%]
Time per replica [ns]
s1
s2
Probability [%]
s1
s2
Conformer

Figure S3: Round-trip times for each replica, calculated from the REST-RECT simulations employing the CHARMM36m force field. The x-axis represents the progression over time, plotted against the duration of each round trip in ns. Every blue cross indicates the duration of one round trip, whereas the red dotted line the average over all recorded round trips in that replica. Calculations only include the simulation using starting structure s1 for simplicity, respectively.

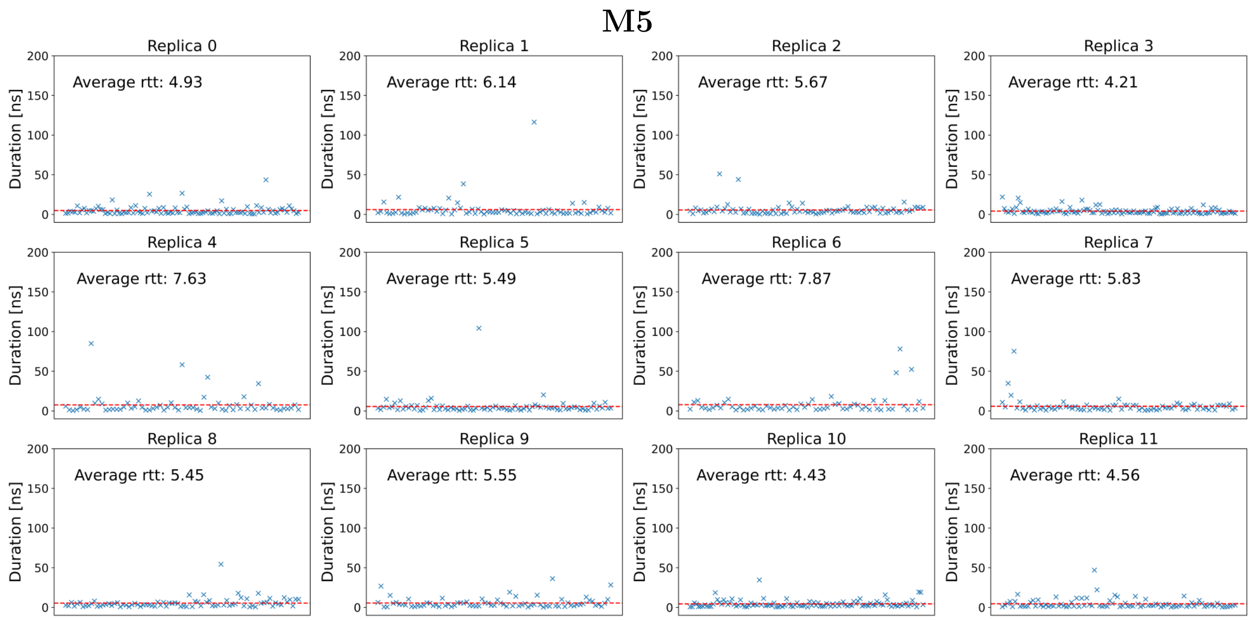


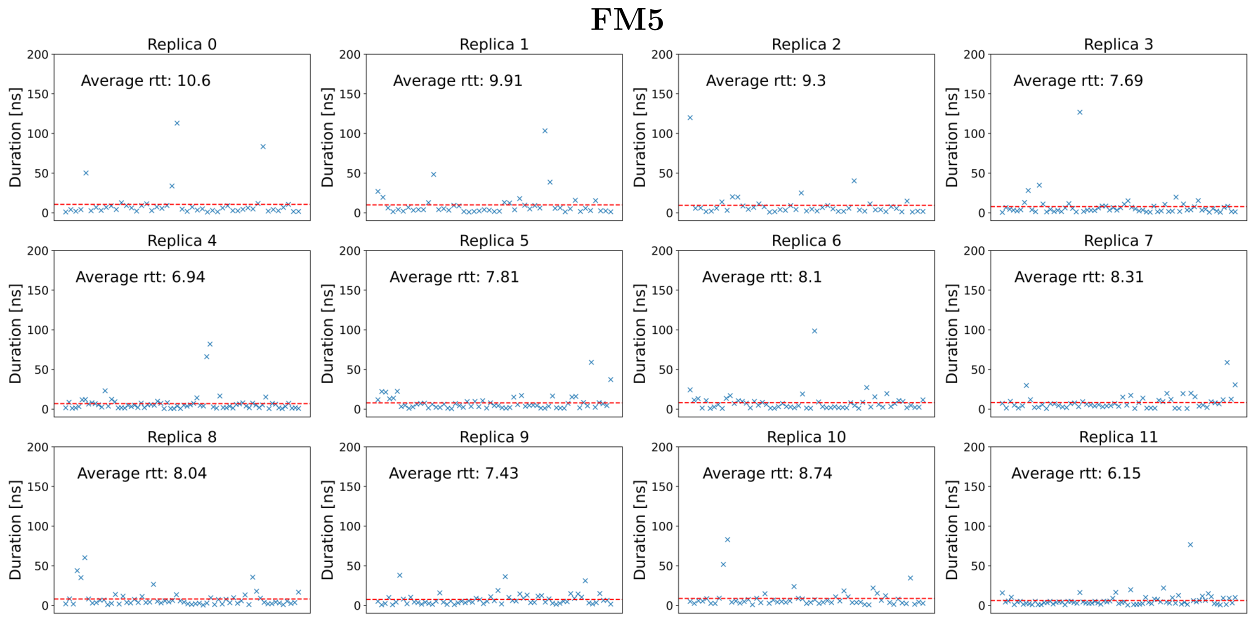

# M9

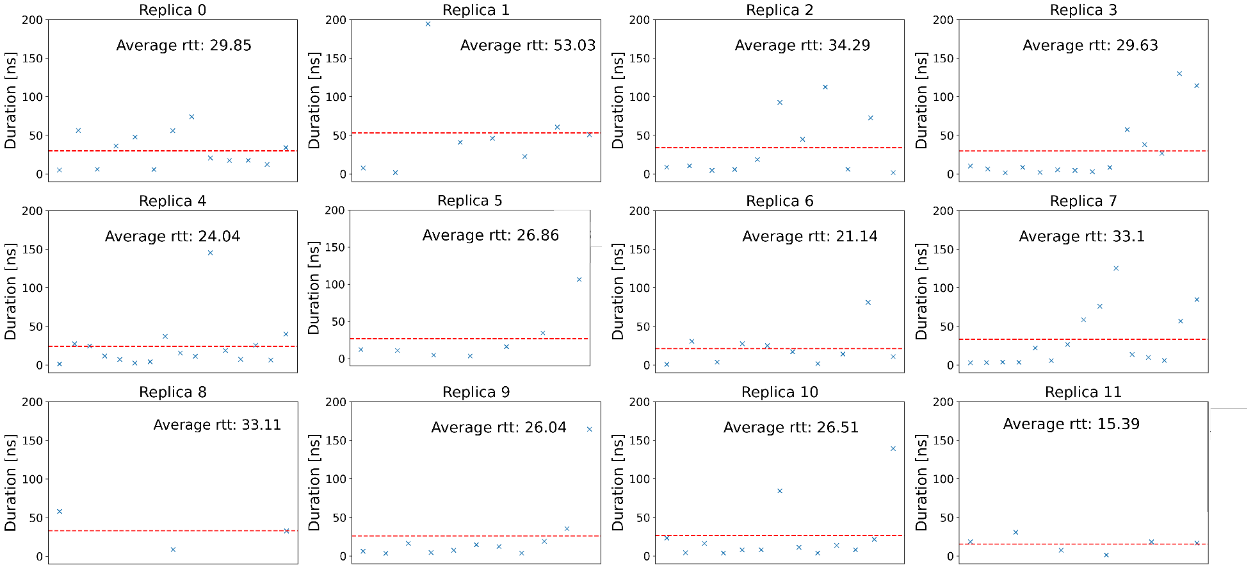
Replica 0
Average rtt: 29.85
Replica 1
Average rtt: 53.03
Replica 2
Average rtt: 34.29
Replica 3
Average rtt: 29.63
Replica 4
Average rtt: 24.04
Replica 5
Average rtt: 26.86
Replica 6
Average rtt: 21.14
Replica 7
Average rtt: 33.1
Replica 8
Average rtt: 33.11
Replica 9
Average rtt: 26.04
Replica 10
Average rtt: 26.51
Replica 11
Average rtt: 15.39
Duration [ns]

# A2G2S2

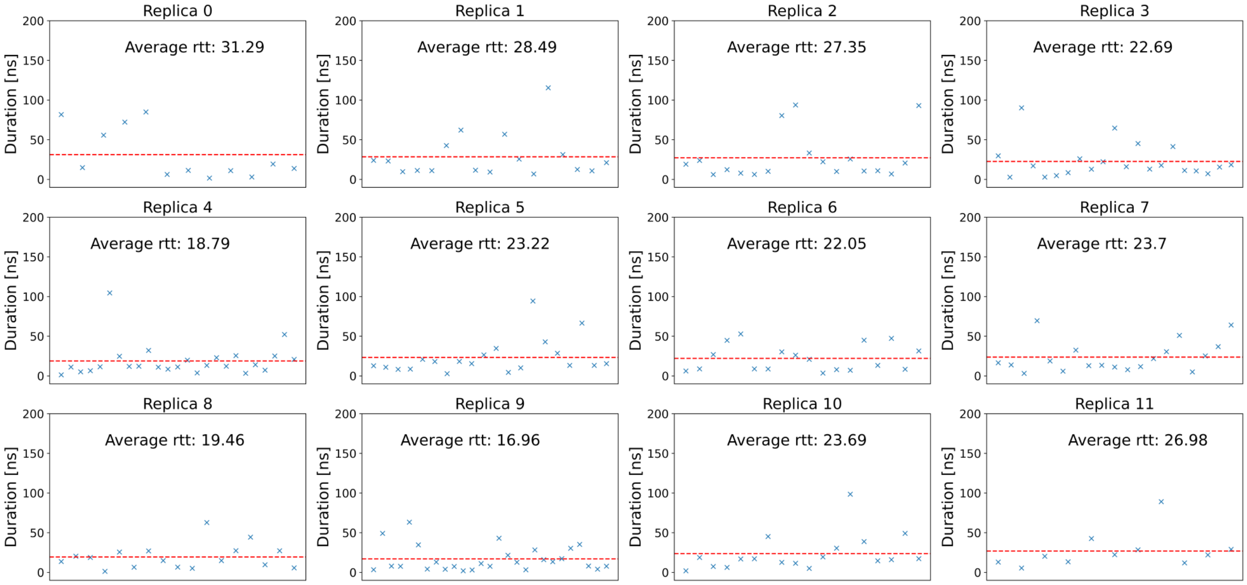
Replica 0
Average rtt: 31.29
Replica 1
Average rtt: 28.49
Replica 2
Average rtt: 27.35
Replica 3
Average rtt: 22.69
Replica 4
Average rtt: 18.79
Replica 5
Average rtt: 23.22
Replica 6
Average rtt: 22.05
Replica 7
Average rtt: 23.7
Replica 8
Average rtt: 19.46
Replica 9
Average rtt: 16.96
Replica 10
Average rtt: 23.69
Replica 11
Average rtt: 26.98
Duration [ns]

# A2G2

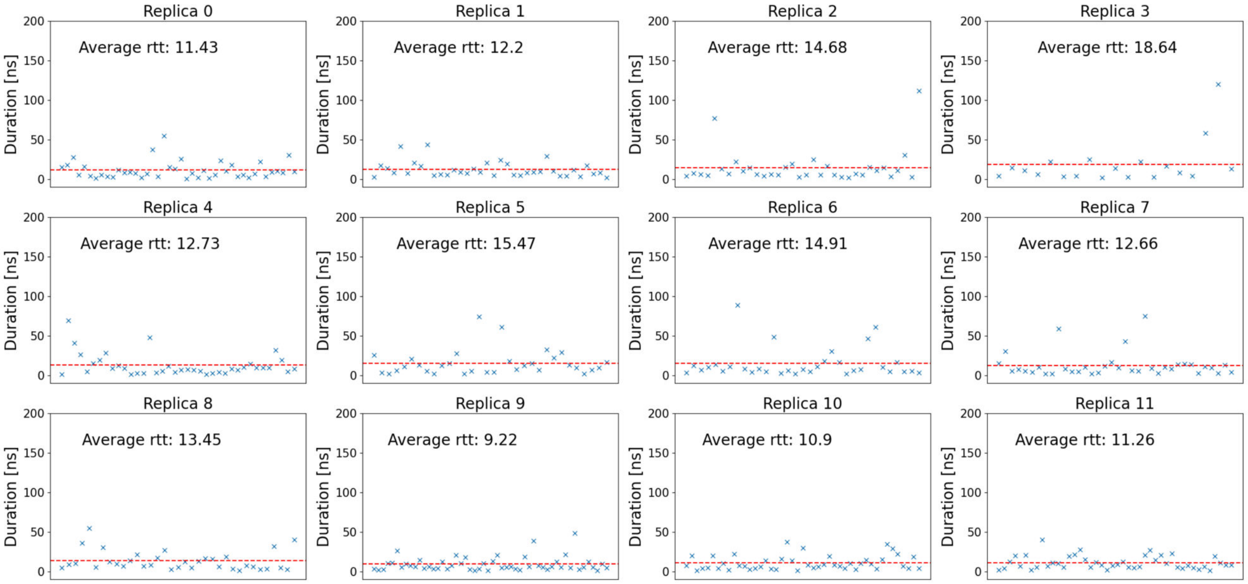

Replica 0
Average rtt: 11.43
Replica 1
Average rtt: 12.2
Replica 2
Average rtt: 14.68
Replica 3
Average rtt: 18.64
Replica 4
Average rtt: 12.73
Replica 5
Average rtt: 15.47
Replica 6
Average rtt: 14.91
Replica 7
Average rtt: 12.66
Replica 8
Average rtt: 13.45
Replica 9
Average rtt: 9.22
Replica 10
Average rtt: 10.9
Replica 11
Average rtt: 11.26
Duration [ns]
0
50
100
150
200

#### GLYCAM06j

Figure S4: Comparison of REST-RECT with plain MD simulations for systems M5, M9, A2G2S2 (and A2G2) employing the GLYCAM06j FF. The upper panel includes atomistic structures of each glycan. The middle panel shows the moving average for the three most populated conformers using a window size of 100 ns (REST-RECT) and 1.2 $\mu$s (MD), corresponding to the same sampling time. Two separate simulations were performed with differing initial starting configurations (s1,s2). The lower panel represents conformer distributions for both starting structures. The conformer string is given on the x-axis, where each digit stands for a torsion angle, the letter representing the occupied minima. Dots are used instead of letters when no change in occupancy could be observed in comparison with the most occupied conformer. The gray box represents a difference in conformer digits between REST-RECT and MD simulations.

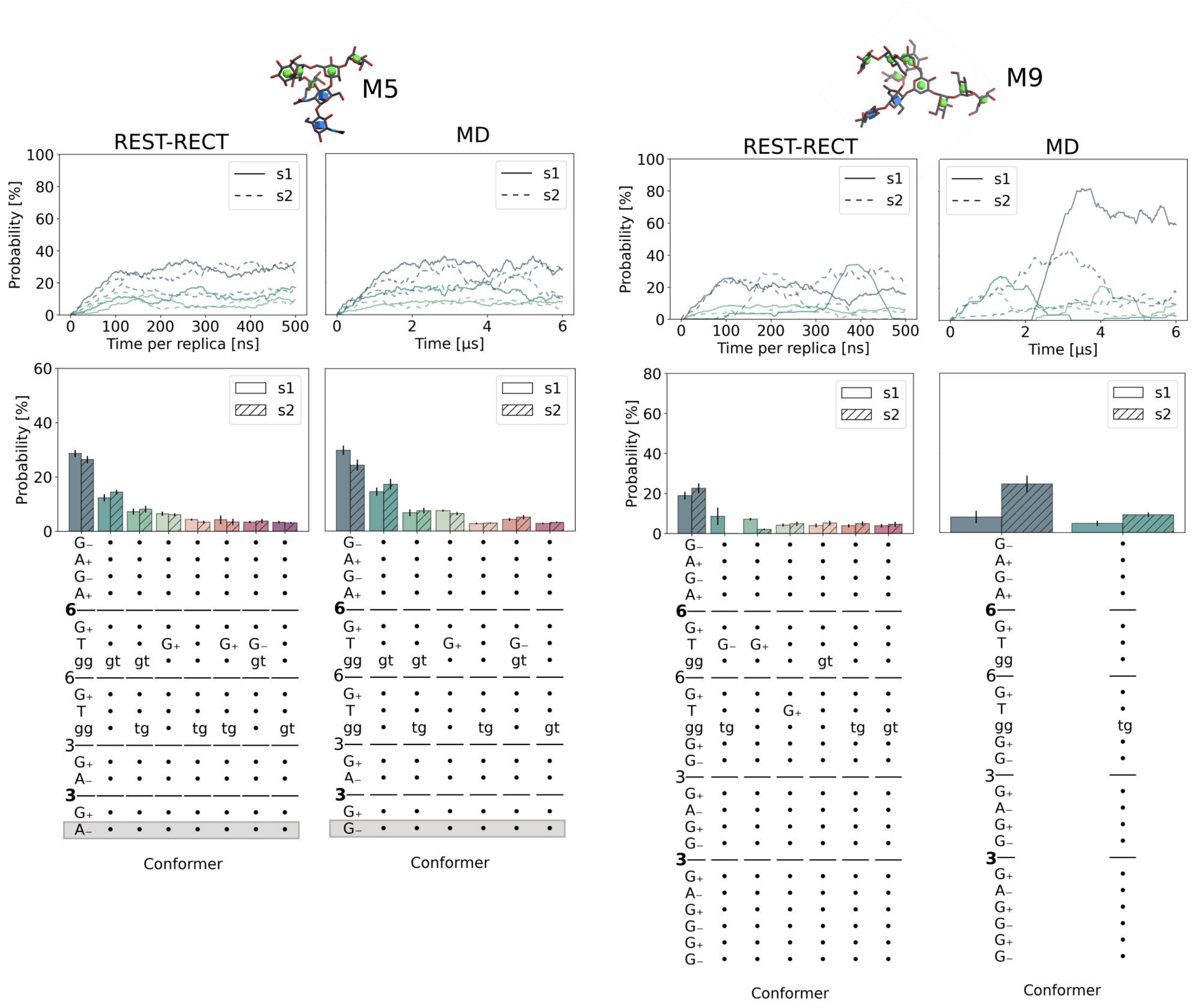

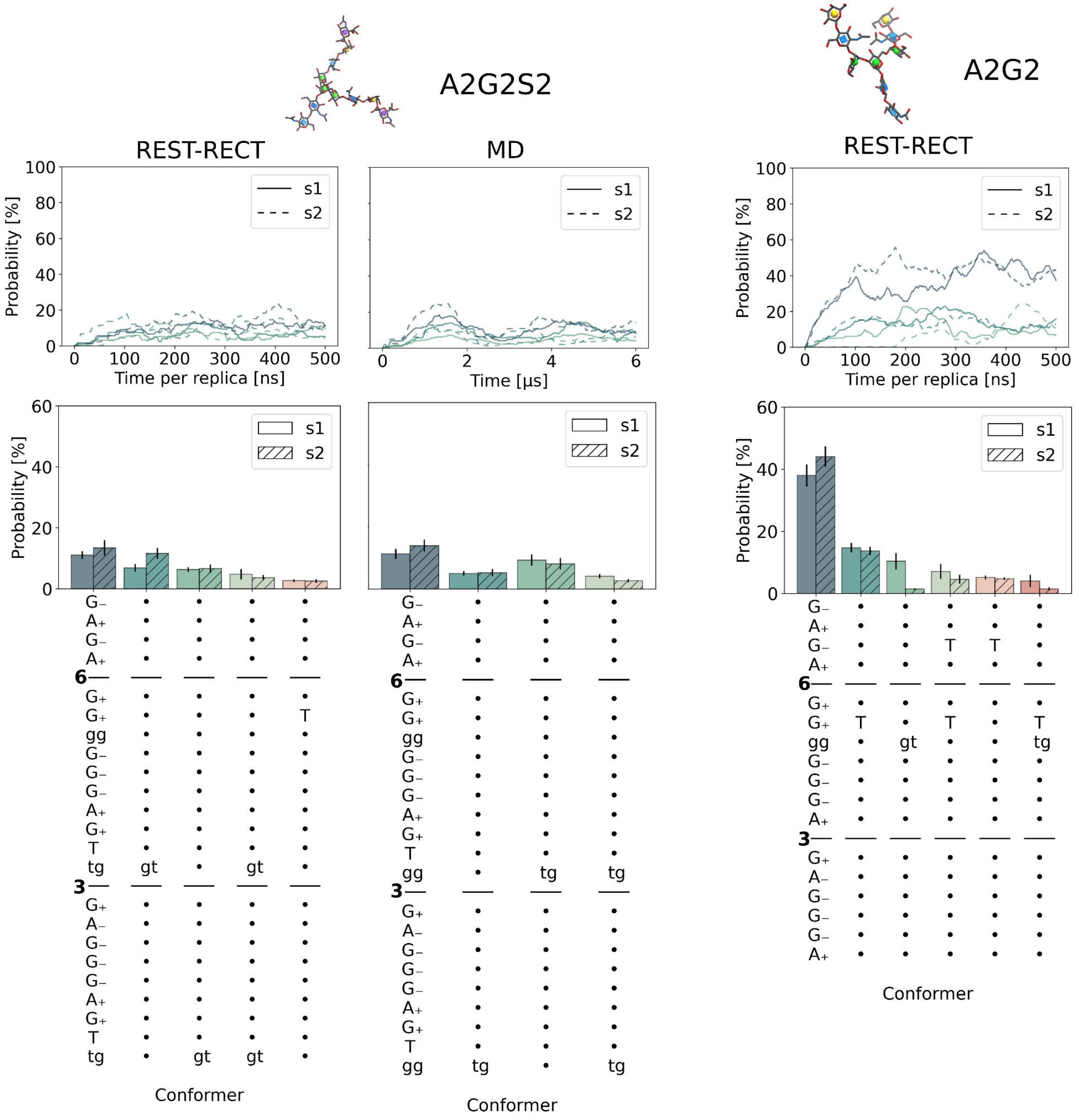

A2G2S2
A2G2
REST-RECT
MD
REST-RECT
Probability [%]
Time per replica [ns]
Time [µs]
s1
s2
Conformer

Figure S5: Rtt for each replica, calculated from the REST-RECT simulations employing the GLYCAM06j FF. The x-axis represents the progression over time, plotted against the duration of each round trip in ns. Every blue cross indicates the duration of one round trip, whereas the red dotted line the average over all recorded round trips in that replica. Calculations only include the simulation using starting structure s1 for simplicity, respectively.

**M5**

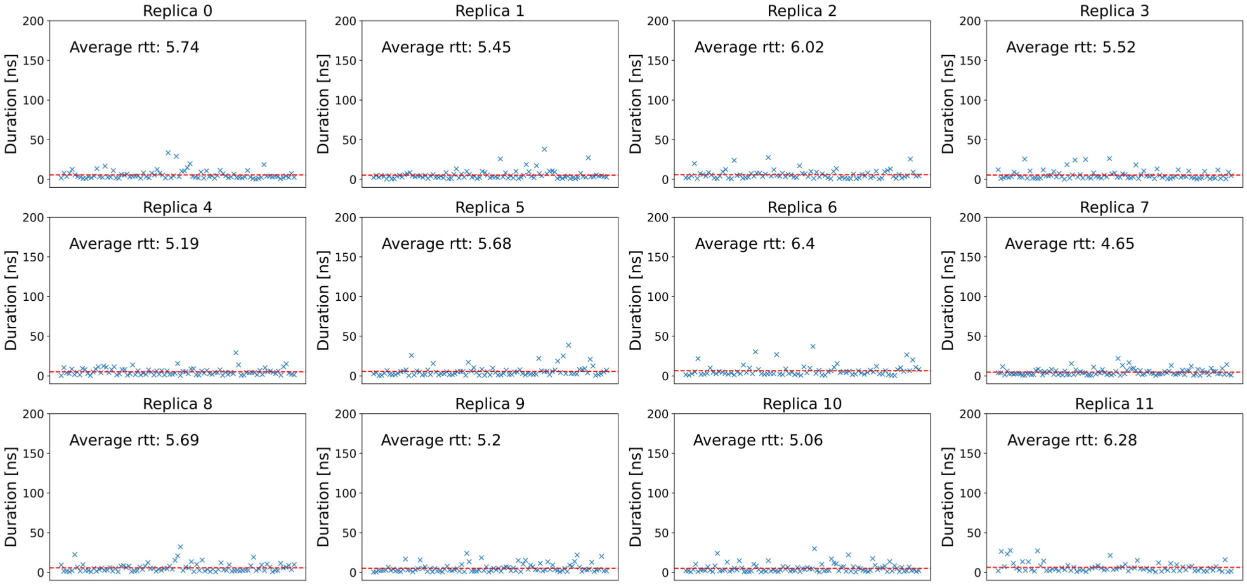


**M9**

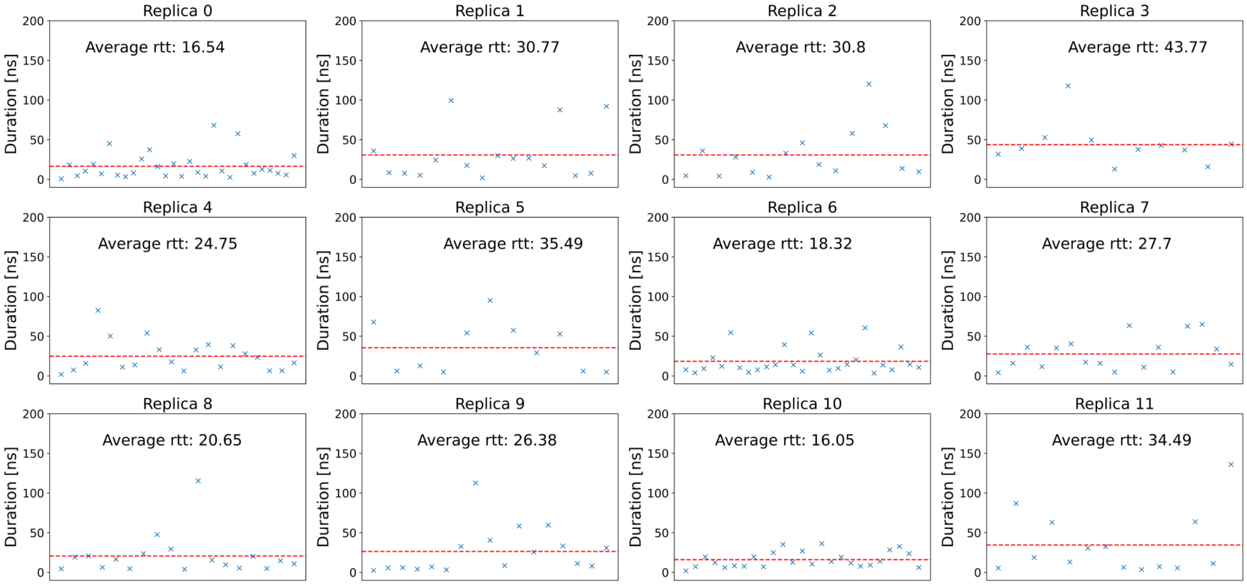

# A2G2S2

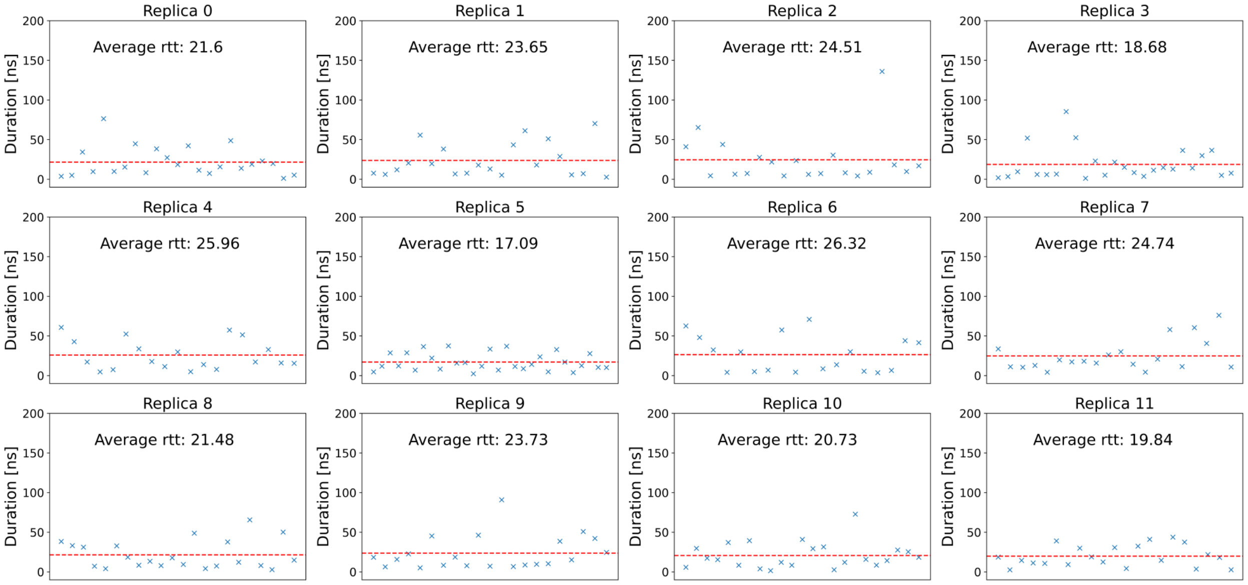

Replica 0
Average rtt: 21.6
Replica 1
Average rtt: 23.65
Replica 2
Average rtt: 24.51
Replica 3
Average rtt: 18.68
Replica 4
Average rtt: 25.96
Replica 5
Average rtt: 17.09
Replica 6
Average rtt: 26.32
Replica 7
Average rtt: 24.74
Replica 8
Average rtt: 21.48
Replica 9
Average rtt: 23.73
Replica 10
Average rtt: 20.73
Replica 11
Average rtt: 19.84
Duration [ns]
0
50
100
150
200


# A2G2

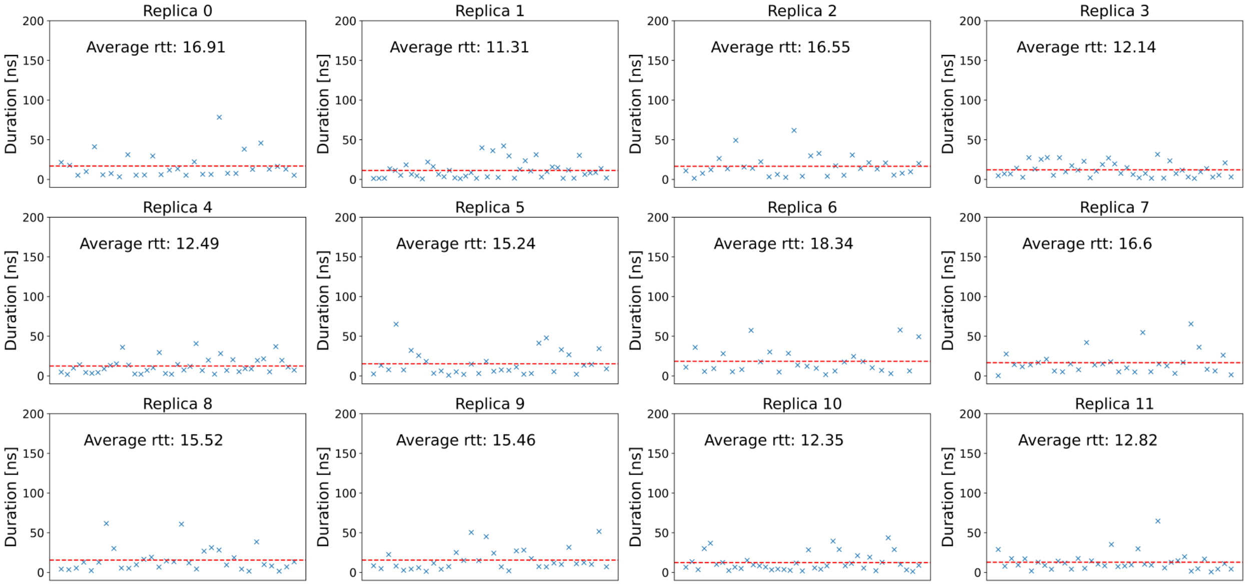

Replica 0
Average rtt: 16.91
Replica 1
Average rtt: 11.31
Replica 2
Average rtt: 16.55
Replica 3
Average rtt: 12.14
Replica 4
Average rtt: 12.49
Replica 5
Average rtt: 15.24
Replica 6
Average rtt: 18.34
Replica 7
Average rtt: 16.6
Replica 8
Average rtt: 15.52
Replica 9
Average rtt: 15.46
Replica 10
Average rtt: 12.35
Replica 11
Average rtt: 12.82
Duration [ns]
0
50
100
150
200

#### Dimensionality reduction of N-glycans

Figure S6: Performance analysis of the four different dimensionality reduction techniques: Principle component analysis (PCA),diffusion map, sketch-map and kernel principle covariates regression (kPcovR). Eigenvalues of the covariant matrix (y-axis) are plotted against the corresponding components (x-axis) for PCA and diffusion maps. For sketch-map, the histogram of pairwise distances plotted for randomly picked points at a given distance ($R_{ij}$) is shown. The parameter $\sigma$ was chosen from this probability distribution, as it represents the switching distance for the sigmoid function, defining which distances are considered close or far in the sketch-map algorithm. At last, regression of the kPcovR algorithm is shown by plotting the property matrix $\mathbf{Y}$ against the reconstructed property variables $\hat{\mathbf{Y}}$. Points are colored according to their associated loss.

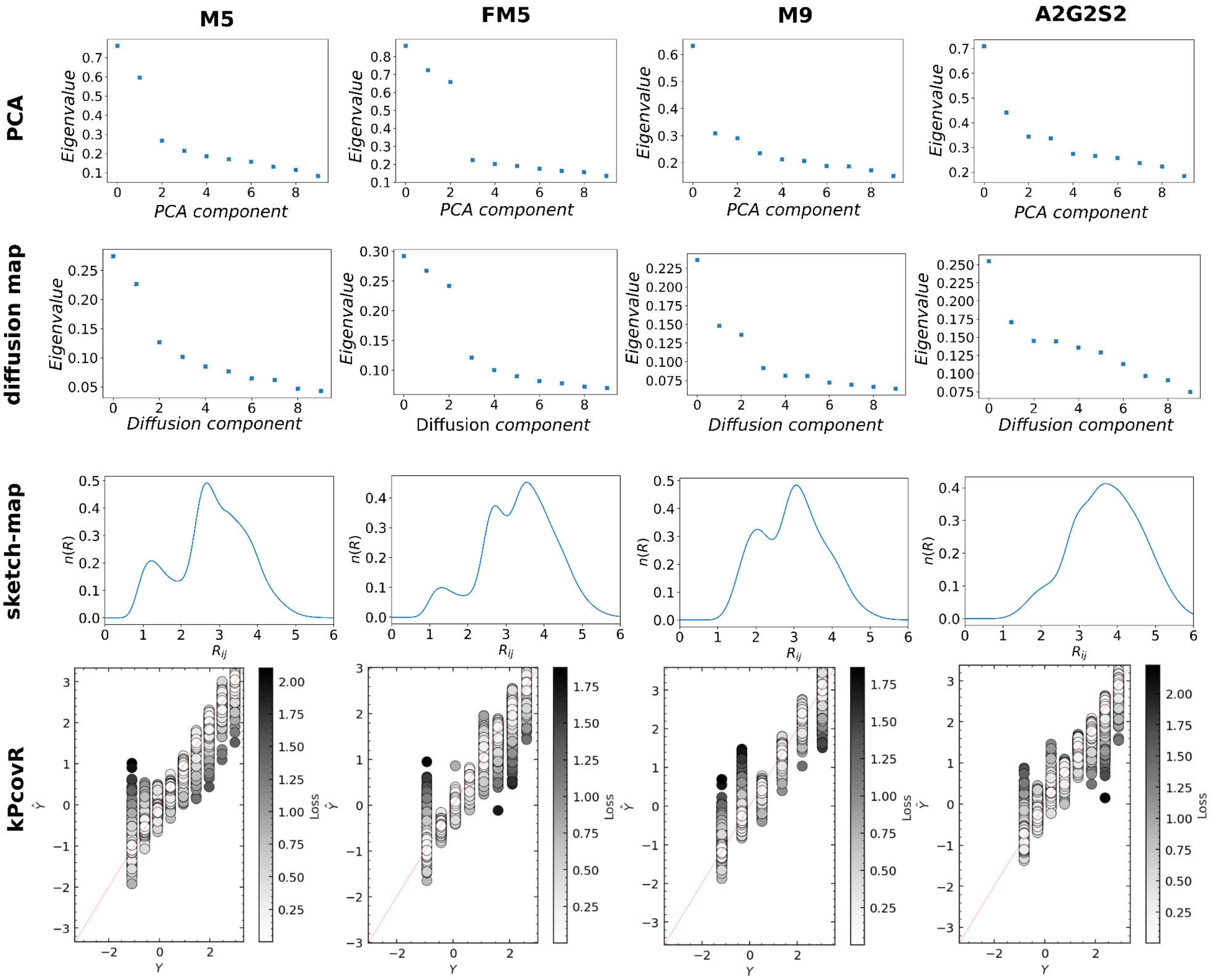

Figure S7: Comparison of four different dimensionality reduction algorithms to cluster the distinct conformers of *N*-glycans FM5, M9 and A2G2S2 apart. PCA, diffusion map and sketch-map employ all torsion angles of each glycan as features, whereas kPcovR additionally uses the conformer strings as properties. Each gray point corresponds to one frame and coloured points to the respective conformers given in Figure S2.

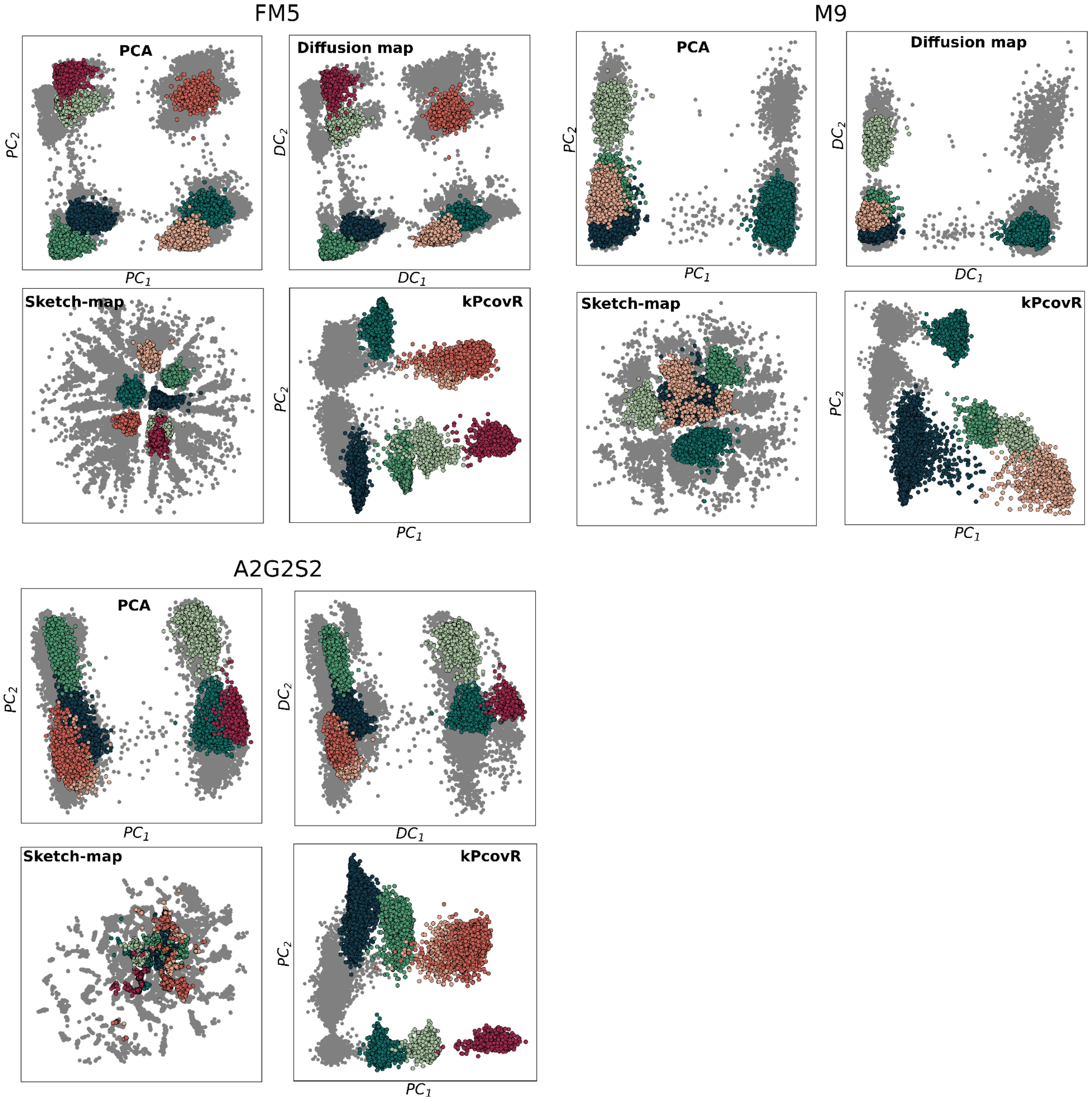

#### Force field comparison

Figure S8: Evaluation of force field performance for M5, M9 and A2G2 examined by PCA and conformer distribution comparison. REST-RECT simulations of each *N*-glycan were performed using either the CHARMM36m or GLYCAM06j FF, initializing simulations from two independent starting conformers (s1,s2). The upper panel represents a PCA of the conformer distribution and its conversion to a free energy surface, respectively for both force fields. Colors in the middle PCA are in accordance to the conformer distributions in the lower panel. Free energy plots along torsion angles (indicated by gray boxes) are represented at the sides with labeled free energy minima. Only those differing substatially between both force fields are displayed.

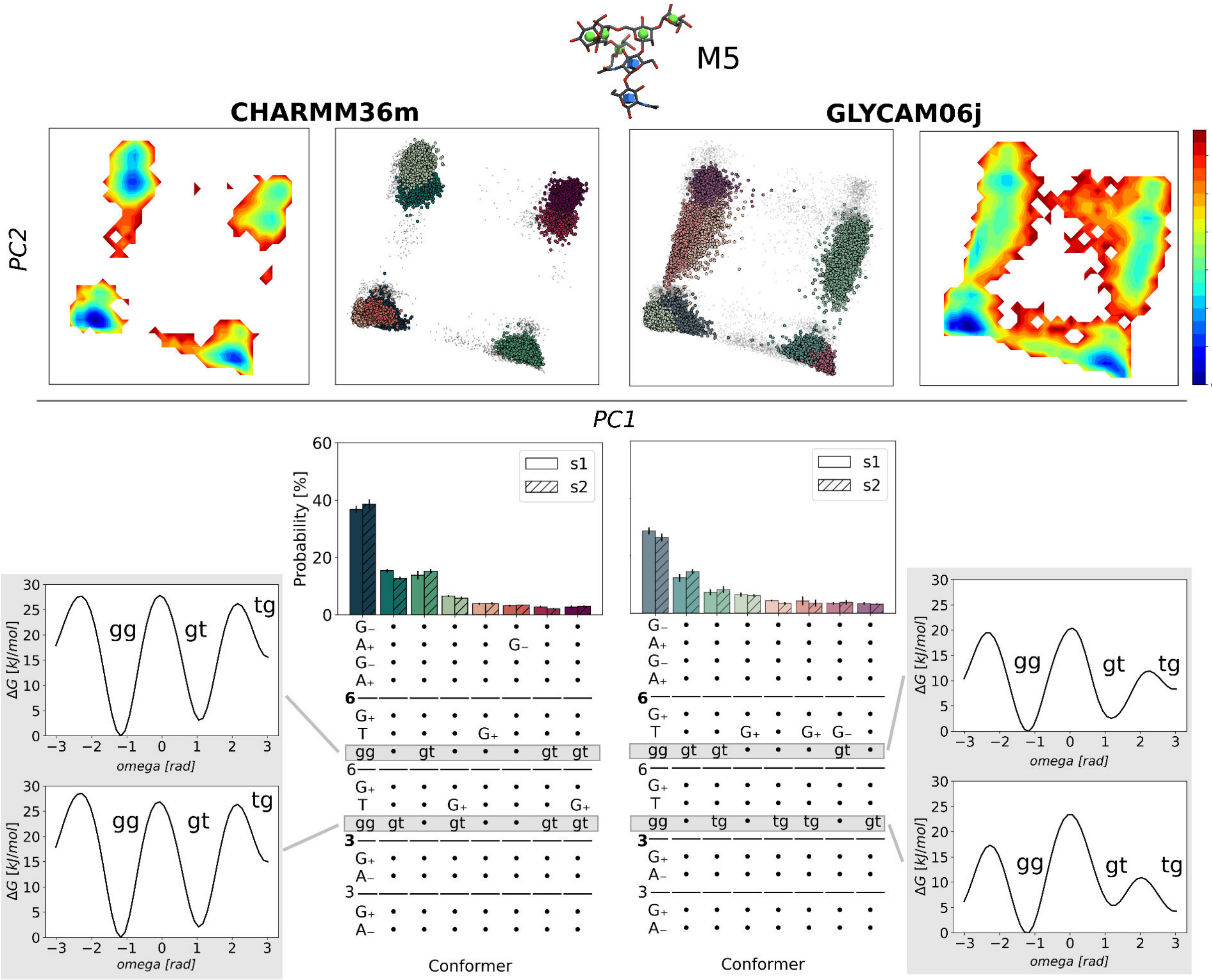

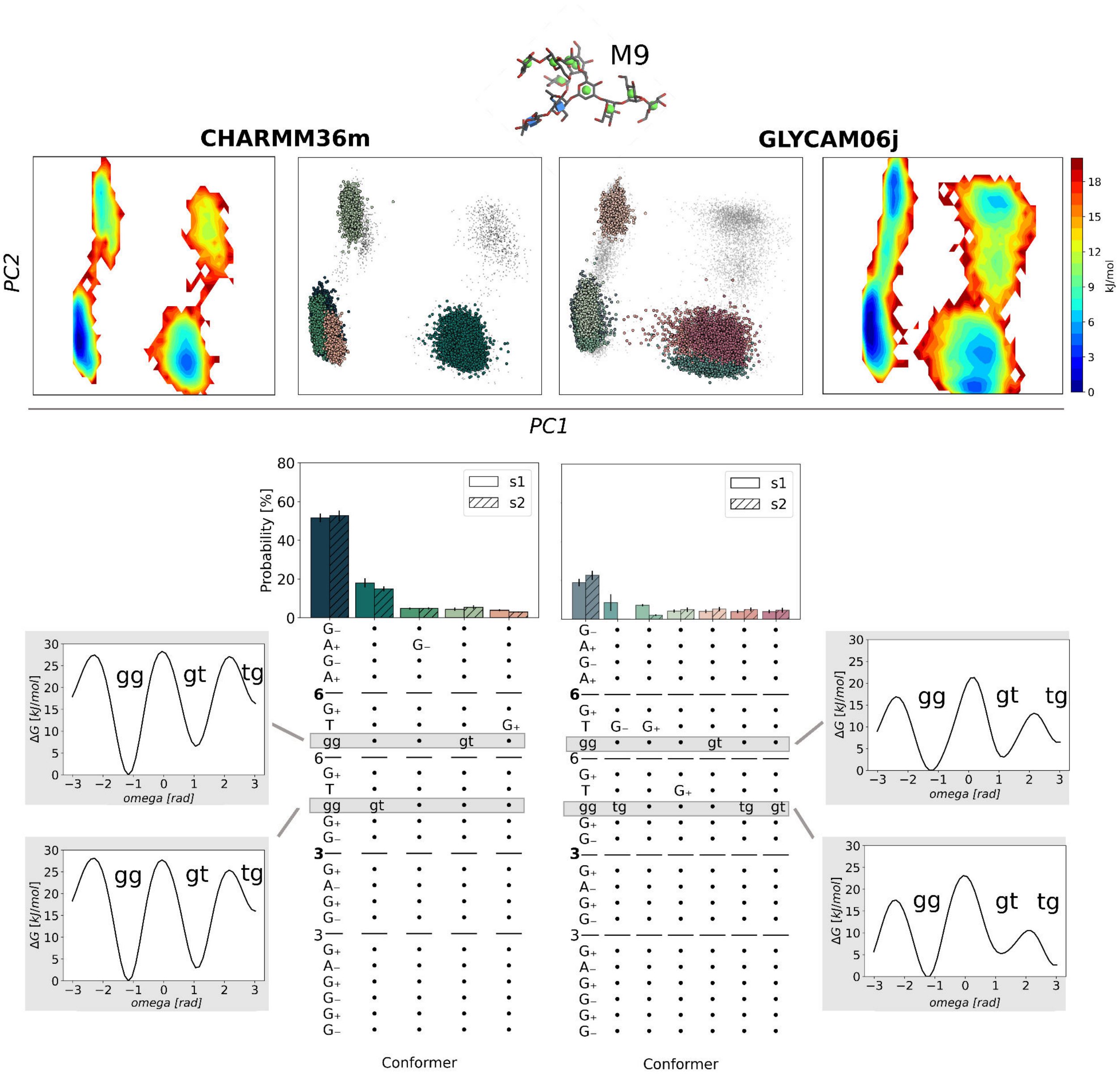
M9
CHARMM36m
GLYCAM06j
PC2
PC1
kJ/mol
Probability [%]
s1
s2
Conformer
Conformer
ΔG [kJ/mol]
omega [rad]
gg
gt
tg

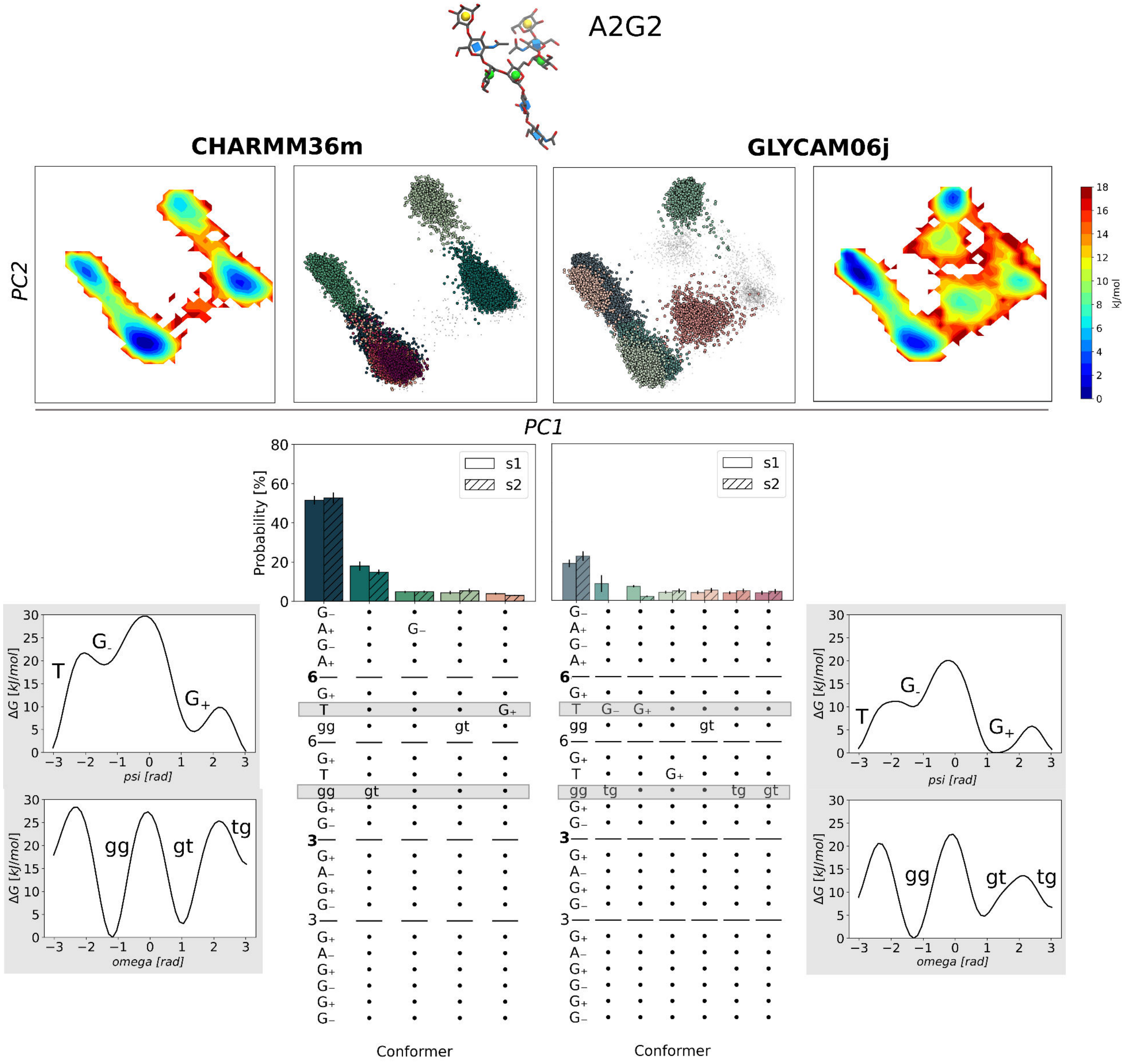
A2G2
CHARMM36m
GLYCAM06j
PC2
PC1
kJ/mol
Probability [%]
s1
s2
ΔG [kJ/mol]
psi [rad]
omega [rad]
T
G-
G+
gg
gt
tg
Conformer
Conformer

#### Free energy profiles

Figure S9: Free energy surfaces along all torsion angles of $N$-glycan A2G2S2, explicitly biased in REST-RECT simulations and constructed by reweighting. Plots of each torsion are ordered according to their occurance in the conformer string, starting from the top left and descending until reaching the bottom and continuing in the next column.

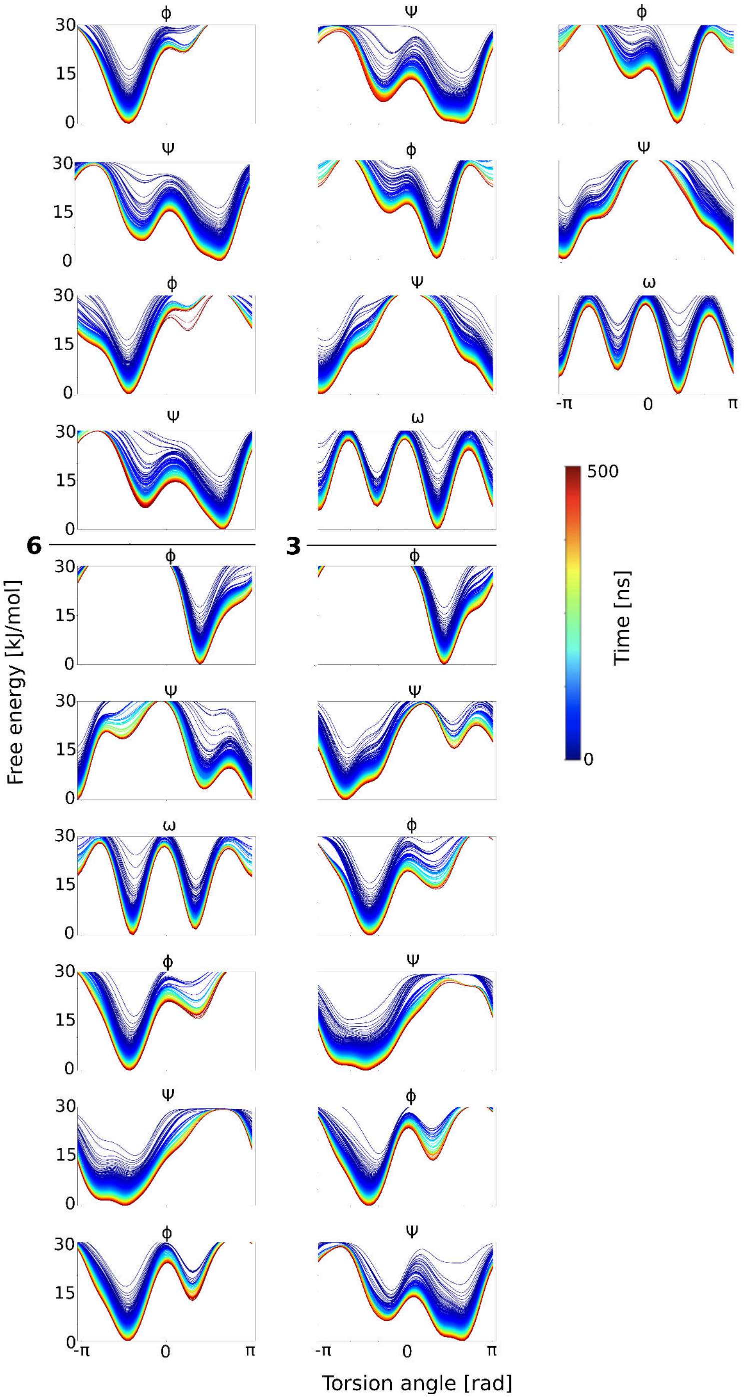

Figure S10: Free energy surfaces along the Cremer-Pople puckering coordinates $\theta$ and $\phi$ for all residues of $N$-glycan A2G2S2, comparing the two force fields CHARMM36m and GLYCAM06j. Collective variables were computed from ground replica REST-RECT simulations by histogram construction and convertion to free energies.

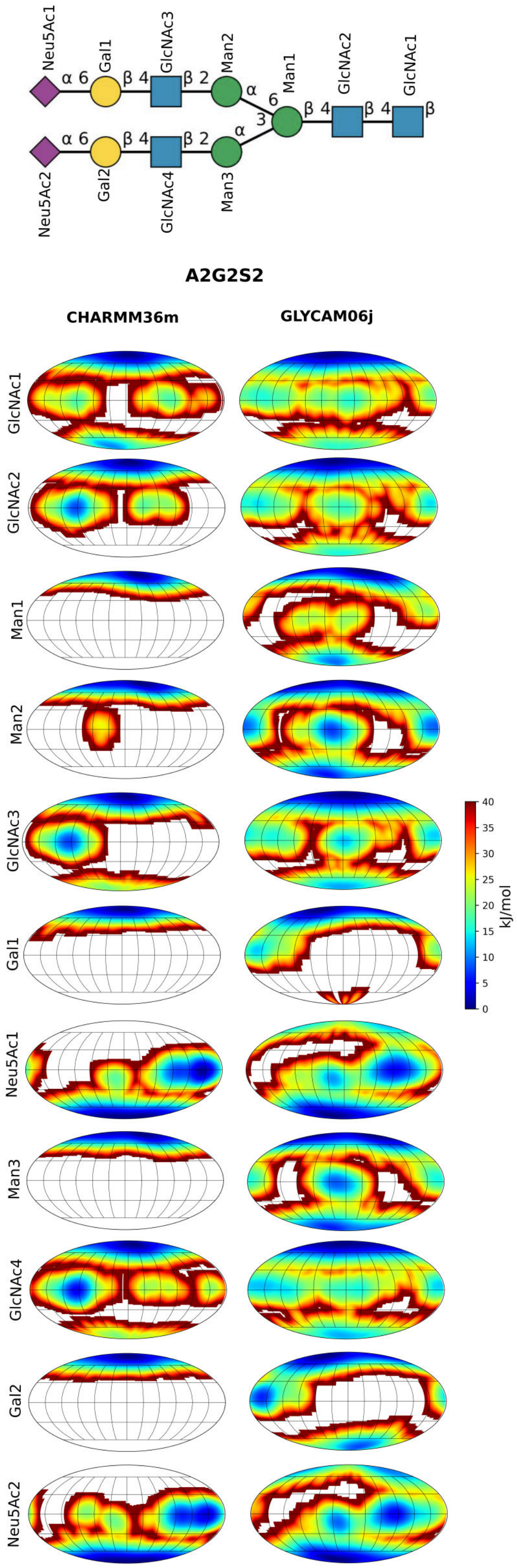

Figure S11: Free energy profile of A2G2, projected in a two dimensional phase along principle components 1 and 2, constructed by PCA. Labeling of conformers into categories is based on the $\psi$ and $\omega$ angle values of the 1-6 linkage.

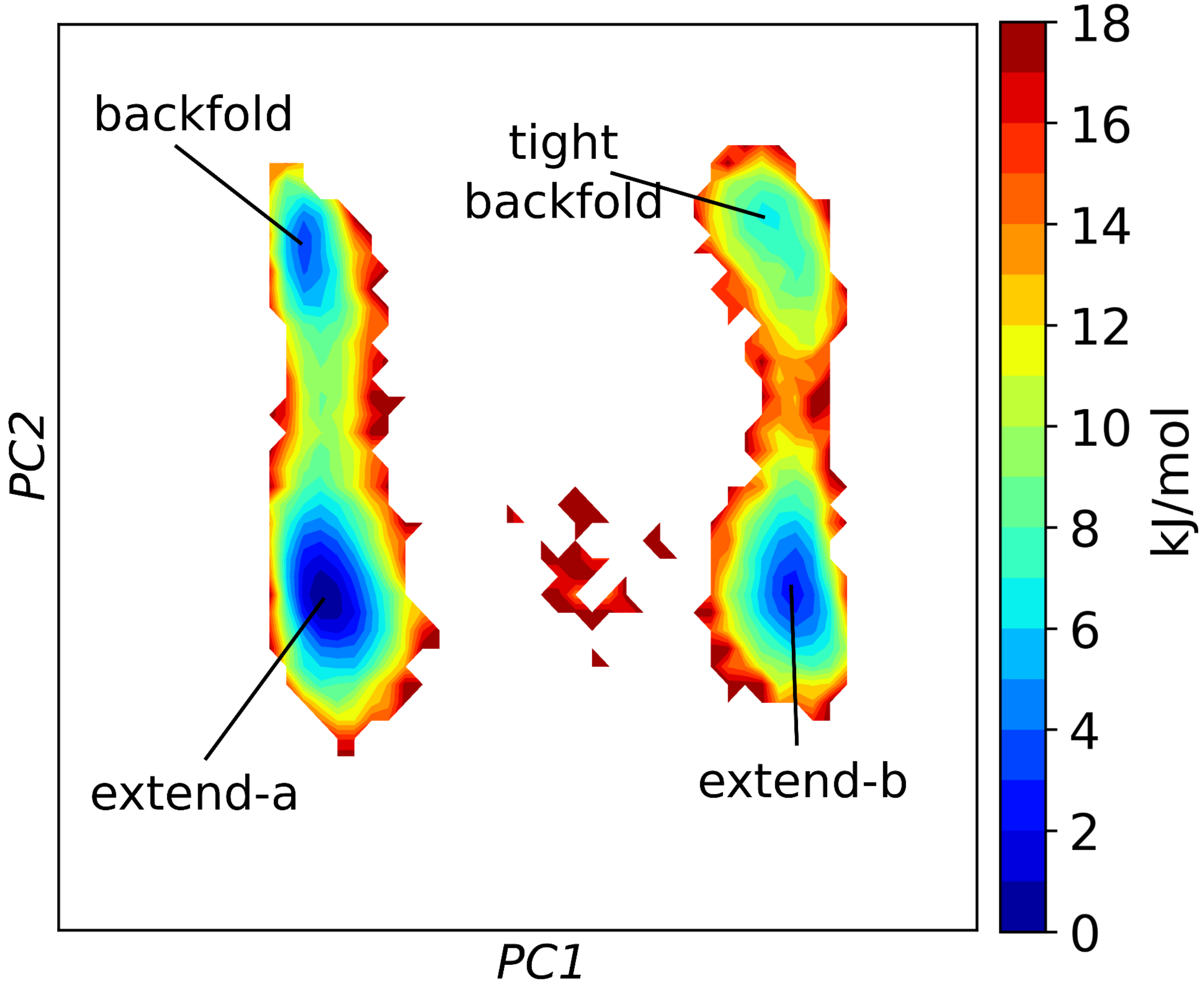


**Force field validation via NMR parameter**

Table S1: Validation of $\omega$ angle populations by NMR J-couplings for M5, M9, A2G2 and A2G2S2. Values correspond to plots in the main manuscript. Values correspond to three different Karplus-equation variants used: Altona & Haasnoot, Stenutz and Tafazzoli, respectively.

**M5**

**6—**

| | J56 | J56' |
|---|---|---|
| Experimental | 2.1 | 5.8 |
| CHARMM36m | 2.78 ± 0.01 | 3.58 ± 0.12 |
| | 1.55 ± 0.01 | 3.58 ± 0.12 |
| | 1.68 ± 0.01 | 3.91 ± 0.12 |
| GLYCAM06j | 2.95 ± 0.05 | 4.29 ± 0.15 |
| | 1.77 ± 0.05 | 4.86 ± 0.15 |
| | 1.89 ± 0.05 | 4.64 ± 0.14 |

6—

| | J56 | J56' |
|---|---|---|
| Experimental | 2.2 | 5.8 |
| CHARMM36m | 2.78 ± 0.01 | 4.28 ± 0.12 |
| | 1.61 ± 0.01 | 4.80 ± 0.12 |
| | 1.74 ± 0.01 | 4.60 ± 0.12 |
| GLYCAM06j | 3.90 ± 0.14 | 3.02 ± 0.10 |
| | 2.77 ± 0.17 | 3.47 ± 0.09 |
| | 2.83 ± 0.16 | 3.20 ± 0.07 |

**M9**

**6—**

| | J56 | J56' |
|---|---|---|
| Experimental | ∽ 2 | ∽ 2 |
| CHARMM36m | 2.74 ± 0.00 | 2.24 ± 0.12 |
| | 1.41 ± 0.01 | 2.72 ± 0.12 |
| | 1.55 ± 0.01 | 2.66 ± 0.12 |
| GLYCAM06j | 3.15 ± 0.10 | 3.44 ± 0.21 |
| | 1.94 ± 0.11 | 4.01 ± 0.22 |
| | 2.05 ± 0.10 | 3.83 ± 0.21 |

6—

| | J56 | J56' |
|---|---|---|
| Experimental | ∽ 2 | ∽ 2 |
| CHARMM36m | 2.75 ± 0.01 | 3.66 ± 0.22 |
| | 1.50 ± 0.01 | 4.18 ± 0.23 |
| | 1.64 ± 0.01 | 4.02 ± 0.22 |
| GLYCAM06j | 4.44 ± 0.43 | 3.41 ± 0.12 |
| | 3.41 ± 0.51 | 3.83 ± 0.08 |
| | 3.44 ± 0.48 | 3.46 ± 0.07 |

**A2G2**

**6—**

| | J56 | J56' |
|---|---|---|
| Experimental | 2.1 | 5.8 |
| CHARMM36m | 2.80 ± 0.01 | 4.12 ± 0.17 |
| | 1.61 ± 0.02 | 4.65 ± 0.17 |
| | 1.74 ± 0.02 | 4.45 ± 0.16 |
| GLYCAM06j | 3.17 ± 0.19 | 3.33 ± 0.27 |
| | 1.91 ± 0.24 | 3.98 ± 0.25 |
| | 2.03 ± 0.23 | 3.81 ± 0.21 |

**A2G2S2**

**6**— $\alpha$2-6

| | J56 | J56' |
|---|---|---|
| Experimental | 3.9 | 8.4 |
| CHARMM36m | 3.60 ± 0.08 | 9.06 ± 0.06 |
| | 2.95 ± 0.09 | 9.56 ± 0.07 |
| | 2.99 ± 0.08 | 8.89 ± 0.06 |
| GLYCAM06j | 6.04 ± 0.18 | 6.80 ± 0.17 |
| | 6.14 ± 0.18 | 6.39 ± 0.19 |
| | 5.21 ± 0.18 | 6.24 ± 0.17 |

**3**— $\alpha$2-6

| | J56 | J56' |
|---|---|---|
| Experimental | 3.9 | 8.4 |
| CHARMM36m | 3.67 ± 0.10 | 9.09 ± 0.13 |
| | 3.01 ± 0.11 | 9.58 ± 0.14 |
| | 3.06 ± 0.10 | 8.91 ± 0.14 |
| GLYCAM06j | 5.89 ± 0.15 | 7.07 ± 0.26 |
| | 5.98 ± 0.18 | 6.69 ± 0.28 |
| | 5.01 ± 0.21 | 6.53 ± 0.27 |
